# Fetal cannabidiol (CBD) exposure alters thermal pain sensitivity, cognition, and prefrontal cortex excitability

**DOI:** 10.1101/2022.12.06.519350

**Authors:** Karli S. Swenson, Luis E. Gomez Wulschner, Victoria M. Hoelscher, Lillian Folts, Kamryn M. Korth, Won Chan Oh, Emily Anne Bates

**Author notes:** Corresponding Author: Emily Anne Bates, PhD, Address: 12800 E 19^th^ Avenue, Aurora, CO 80045.

## Abstract

Thousands of people suffer from nausea with pregnancy each year. Nausea can be alleviated with cannabidiol (CBD), a primary component of cannabis that is widely available. However, is it unknown how fetal CBD exposure affects embryonic development and postnatal outcomes. CBD binds and activates receptors that are important for fetal development and are expressed in the fetal brain, including serotonin receptors (5HT_1A_), voltage-gated potassium (Kv)7 receptors, and the transient potential vanilloid 1 receptor (TRPV1). Excessive activation of each of these receptors during fetal development can disrupt neurodevelopment. Here, we test the hypothesis that intrauterine CBD exposure alters offspring neurodevelopment and postnatal behavior. We show that fetal CBD exposure sensitizes male offspring to thermal pain in a TRPV1 dependent manner. We show that fetal CBD exposure decreases cognitive function in female CBD-exposed offspring. We demonstrate that fetal CBD exposure increases the minimum current required to elicit action potentials and decreases the number of action potentials in female offspring layer 2/3 prefrontal cortex (PFC) pyramidal neurons. Fetal CBD exposure reduces the amplitude of glutamate uncaging-evoked excitatory post-synaptic currents. Combined, these data show that fetal CBD exposure disrupts neurodevelopment and postnatal behavior in a sex-dependent manner.

**One Sentence Summary:** Cannabidiol (CBD) consumption during pregnancy alters offspring behavior and neuronal excitability in a sex dependent manner in mice.

## Introduction

The nausea and vomiting of morning sickness is debilitating for thousands of pregnant patients each year^1^. Pregnant people are drawn to use cannabis for its anti-emetic, or anti-nausea, properties because they believe it to be safe^2^. Cannabis has two primary component parts, cannabidiol (CBD) and tetrahydrocannabinol (THC), along with minor cannabinoids and terpenes. Although research quantifying CBD consumption in a pregnant population is limited, THC metabolites were detected in cord blood samples from twenty-two percent of pregnant people under the age of twenty-five tested in Colorado, suggesting CBD consumption in the same group^3^. CBD is an effective anti-emetic medication, but does not induce the psychoactive properties of THC^4^. CBD has become widely available since it was removed from schedule 1 drug classification in 2018^5^. In addition to the number of pregnant people who consume CBD as a component of whole cannabis, it is a likely that a population of pregnant patients consume CBD alone^6^ due to the availability of CBD, and its anti-emetic properties.

CBD diffuses through maternal-placental-fetal circulation^6^. Animal models show that lipophilic CBD accumulates in the fetal brain, liver, and gastrointestinal tract^6^. CBD binds and activates receptors important for fetal brain development including the 5HT_1A_ serotonin receptor, heat-activated transient potential vanilloid receptor one (TRPV1) calcium channels^7^, and voltage-gated Kv7 receptor potassium channel^8,9,10^, among others.

Excessive TRPV1 activation in neural crest cells confers neural tube defects akin to those induced by maternal fever^11^, suggesting that increased activation of TRPV1 affects developmental processes. Fetal overactivation of TRPV1 induces anxiety behaviors in mice^12^. Excessive activation of TRPV1 alters excitatory innervation in hippocampal neurons and mediates action potential threshold^13^. CBD also activates TRPV2, a calcium channel with a higher temperature threshold than TRPV1^14^. Excessive activation of TRPV2 during development alters axon extension and the release of calcitonin gene related peptide (CGRP), a proinflammatory signaling molecule, in the dorsal root ganglion^14^. Fetal cannabis exposure is associated with increased anxiety in humans, but it is unknown whether CBD contributes to this association^15^. TRPV1 is expressed in the hypothalamus and the rostral hindbrain in rat, primate, and human tissue^16^. TRPV1 is expressed in cortical layers 3 and 5 of the hippocampus, central amygdala, habenula, striatum, hypothalamus, centromedial, and paraventricular thalamic nuclei, substantia nigra, reticular formation, locus coeruleus, cerebellum, and inferior olive in adult rodent brains^17^ and in the dorsomedial hypothalamus, hippocampus, dorsal root ganglion cells, supramammillary nucleus, and periaqueductal grey in fetal mice^16^. These limbic regions mediate behavioral and emotional responses to stimuli^18^. Because CBD activates TRPV1 and TRPV2, and these calcium channels are expressed in the fetal brain, and their excessive activation during fetal development is associated with negative developmental effects, we hypothesize that fetal CBD exposure could disrupt brain development and postnatal behavior.

Excessive fetal serotonin signaling harms neuronal development. Overexpression of 5HT_1A_ during mouse fetal and early postnatal development decreases adult anxiety behaviors on elevated plus maze test and free exploratory paradigm, and decreases spatial learning via the Morris water maze compared to wild type mice^19^. Excessive activation of 5HT_1A_ during fetal development decreases neurogenesis, decreases neuron network complexity, alters neuron refinement, delays sensory-evoked potentials, decreases sensory evoked firing, and decreases amplitude of sensory evoked potentials^20^. Depletion of tryptophan, the molecular precursor to serotonin, impairs cognition in humans and mice^21,22^. 5HT receptor transcripts are expressed from embryonic day (E)14.5-16.5 in the thalamus, hippocampus, and in a medial to lateral gradient in the cortex in mice^23^. Human fetal brains have highest expression of 5HT_1A_ between the sixteenth and twenty-second weeks of gestation in the cortex and the hippocampus^24^. CBD activates 5HT_1A_, and excessive activation is associated with negative outcomes in the fetal brain. We hypothesized that fetal CBD exposure could disrupt similar processes through 5HT_1A_ activation.

CBD binds and activates Kv7.2 and Kv7.3, which are expressed in the brain throughout embryonic and postnatal development^25^. Kv7.2 and Kv7.3 gain-of-function mutations are associated with increased rates of human intellectual disability and epileptic encephalopathy^26^. Kv7.2/3 agonism decreases relative refractory periods and increases post-conditioned super-excitability of neurons in cultured human myelinated axons^27^. Alterations in mouse Kv7.2 and 7.3 activity during development are associated with lasting developmental disorders including intellectual disabilities, memory and behavioral deficits, and disruption of GABAergic inhibitory neuron in the hippocampus humans and neuronal cell culture, respectively^28^. Antagonism of Kv7 channels is under current investigation for potential in improving cognition and memory^29^. CBD activates Kv7 receptors which are expressed in the fetal brain and excessive activation is associated with negative neurodevelopmental effects. We hypothesize that fetal CBD exposure could excessively activate Kv7.2 and Kv7.3 to disrupt brain development and postnatal behavior.

CBD is commonly orally consumed^30^. We administered 50mg/kg of CBD dissolved in sunflower oil or sunflower oil alone to C57BL6 female dams daily from E5 through birth. We selected 50mg/kg in accordance with the national institutes of drugs of abuse (NIDA) recommendation for 5mg/kg via intraperitoneal injection for cannabis research multiplied by 10 to account for the first pass metabolism in the dam liver. We selected oral gavage for a route of administration to model oral CBD consumption with accurately measured daily doses. Offspring were subjected to a battery of behavioral testing from puberty to adulthood to determine if fetal CBD exposure alters postnatal behavior.

We show that fetal CBD exposure disrupts neurodevelopment and postnatal behavior. We show that CBD-exposed male offspring are sensitized to thermal pain in a TRPV1-dependent manner. We found fetal CBD exposure did not affect offspring anxiety, spatial memory, or compulsivity. We show fetal CBD exposure reduces cognition in female mice. We demonstrate that fetal CBD exposure reduces excitability of pyramidal neurons in the prefrontal cortex in postnatal (P)14-21 female mice. Female CBD-exposed offspring require larger currents and more depolarized voltage to elicit action potentials and elicit fewer action potentials at a given current. Fetal CBD exposure reduced the amplitude of glutamate uncaging-evoked excitatory post-synaptic currents in female prefrontal cortical slices. CBD metabolites were not retained in pup plasma by P8 suggesting that fetal CBD exposure changes fetal neuronal physiology and neurodevelopment to impact postnatal behavior.

## Results

### CBD metabolites are present in dam, fetal, and neonatal blood plasma

We administered 50mg/kg CBD dissolved in sunflower oil or sunflower oil alone (vehicle) via oral gavage daily from E5 through birth to C57BL6J or *TRPV1*^*KO/KO*^ female mice (Figure 1A). CBD is metabolized into 6a-hydroxy cannabidiol, 7-hydroxy cannabidiol, carboxy-cannabidiol, and cannabidiol glucuronide and each crosses the placenta. We used liquid chromatography tandem mass spectrometry to quantify CBD and its metabolites in dam and pup plasma from two hours after dosing on E18.5 and postnatal day (P) 0, P4, P8, and P12. We found 630.93 ± 383.72 ug/ml CBD, N=3 dams at E18.5, 250.24 ± 130.51 ug/ml CBD, N=4 at P0, and 2.05 ug/ml ± 1.52 CBD, N=3 at P4 in dam plasma. We found 588.57 ± 253.32 ug/ml CBD, N=3 pooled litters at E18.5, 139.73 ± 98.39 ug/ml CBD, N=6 pooled litters at P0, and 0.83 ± 0.53 ug/ml CBD, N=4 pooled litters at P4 (Figure 1B). Additional CBD metabolite levels for 6a-hydroxy cannabidiol, 7-hydroxy cannabidiol, carboxy-cannabidiol, and cannabidiol glucuronide can be found in supplemental table 1. By P8, dams and pups had fully cleared the CBD and its metabolites were below the limit of detection in the plasma (Figure 1B), suggesting that any behavioral or physiological differences between CBD and vehicle-exposed offspring are due to changes in development rather than acute effects of CBD.

**Figure 1.**
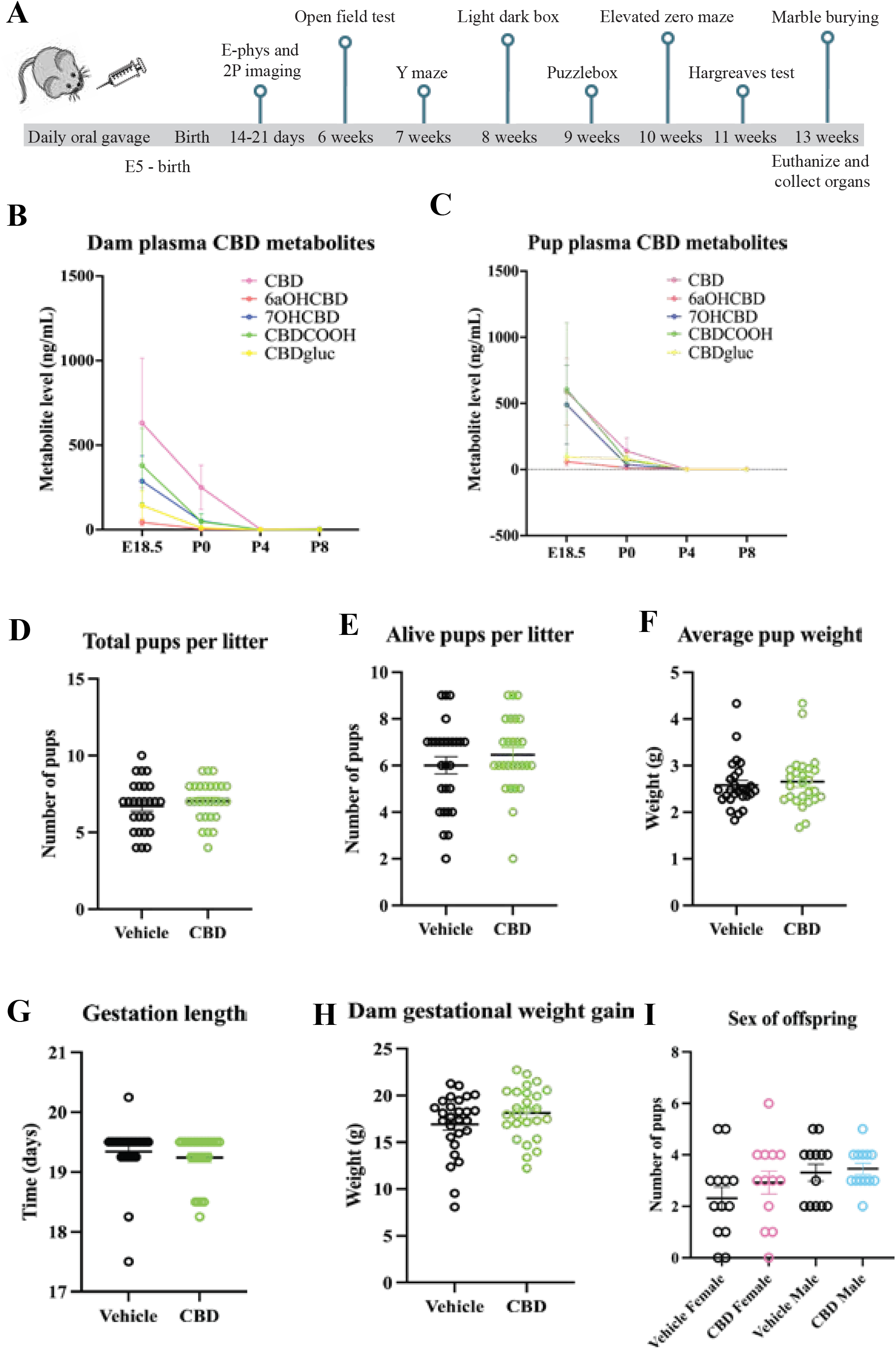
Dosing schematic, validation of CBD metabolites and litter factors. A timeline shows CBD administration and age of offspring when tests were performed (A). A graph shows CBD and CBD metabolites in the dam blood plasma from E18.5, P0, P4, and P8 (B) and pooled pup litter plasma from each group (C) from E18.5, P0, P4, and P8. Graphs show gestational CBD consumption does not alter total pups per litter (D), alive pups per litter (E), average pup weight (F), gestation length (G), gestation weight gain (H), or sex of offspring (I) from 27 vehicle administered dams and 26 CBD administered dams. Error bars represent S.E.M. The gestation length was 19.3 ± 0.092 days (vehicle dams), 19.27 ± 0.073 days (CBD dams), P=0.64, t-test. Gestational weight gain was 16.93 ± 0.6 grams (vehicle dams), 18.11 ± 2.77 grams (CBD dams), P=0.413, t-test. Total pups per litter was 6.70 ± 0.33 (vehicle litters), 7.037 ± 0.25 (CBD litters), P=0.61, t-test. Alive pups per litter was 6.00 ± 0.36 (vehicle litters), 6.44 ± 0.31 (CBD litters), P=0.36, t-test. Vehicle dams birthed 2.31 ± 0.44 female offspring and 3.31 ± 0.33 male offspring (N=13 litters), CBD dams birthed 2.923 ± 0.445 female offspring and 3.46 ± 0.22 male offspring, P=0.34 and P=0.70, respectively, t-tests.

### CBD exposure does not alter pregnancy length, gestational weight gain, litter size, or sex of offspring

To determine how CBD consumption during pregnancy affects maternal factors, we quantified number of pups per litter, pup survival, average pup weight, pup sex ratios, gestation length and dam gestational weight gain in vehicle and CBD dosed dams and their litters. We found that CBD exposure did not alter any of these factors (Figure 1D-I). We conclude that CBD consumption during pregnancy, from E5 through birth, does not induce detectable changes in these maternal factors or litter composition compared to vehicle.

### Fetal CBD exposure increases thermal pain sensitivity in male, but not female, offspring

CBD activates TRPV1, a heat-gated calcium channel^31^. We hypothesized that fetal CBD exposure would excessively activate TRPV1 channels and alter the development of thermal pain circuits. We used the Hargreaves test to measure thermal pain sensitivity in wild type and *TRPV1*^*KO/KO*^ vehicle or CBD-exposed offspring. The Hargreaves test measures the latency to response to a thermal stimulus. Fetal CBD exposure did not affect female sensitivity to thermal pain (12.25 ± 1.46 seconds, N=11 vehicle-exposed versus 14.14 ± 1.21 seconds, N=11 CBD-exposed, P=0.331, t-test). *TRPV1*^*KO/KO*^ vehicle and CBD-exposed females were similarly sensitive to thermal stimuli (16.76 ± 0.60 seconds, N=7 vehicle-exposed *TRPV1*^*KO/KO*^, 14.98 ± 0.82 seconds N=8 CBD-exposed *TRPV1*^*KO/KO*^, P=0.11, t-test. Figure 2A). Female pain tolerance varies with estrus cycle stage^32^. We repeated the Hargreaves test with female offspring controlling for estrus cycle stage and found no differences in thermal pain sensitivity based on fetal CBD exposure (Figure 2A (9.62 ± 1.13 seconds N=9 vehicle-exposed estrus compared to 7.79 ± 0.93 seconds, N=9 CBD-exposed estrus females, P=0.23, t-test). Fetal CBD exposure also did not affect thermal sensitivity in female mice that were not in estrus (11.11 ± 0.78 seconds N=11 vehicle-exposed females 10.99 ± 1.18 seconds N=7 CBD-exposed female mice, P=0.93, t-test). Interestingly, CBD-exposed male offspring were significantly more sensitive to thermal pain (11.58 ± 0.64 seconds, N=8 vehicle-exposed vs 6.87 ± 3.27 seconds, N=8 CBD-exposed, P=4.99E-8, t-test). The effect of CBD on thermal sensitivity in males is dependent on the TRPV1 receptor. CBD exposure did not impact thermal sensitivity in *TRPV1*^*KO/KO*^ male offspring (11.09 ± 0.65 seconds, N=8 vehicle-exposed, vs. 12.43 ± 1.61 seconds, N=8, P=0.45, t-test, Figure 2A). These data show that oral consumption of 50 mg/kg CBD during pregnancy was sufficient to increase thermal pain sensitivity in adult male offspring.

**Figure 2.**
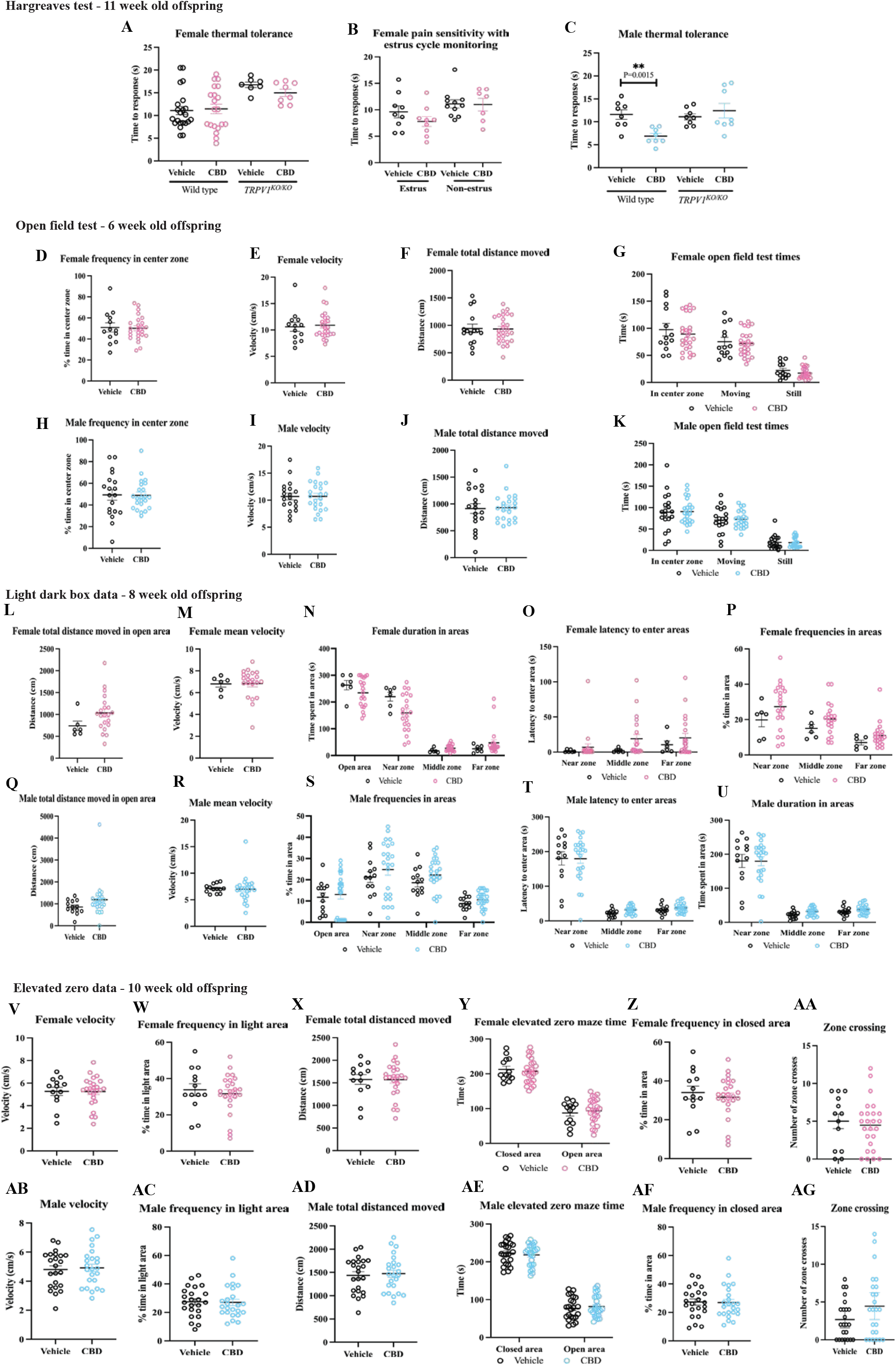
Fetal CBD exposure increases thermal sensitivity in male mice, but not female mice. Fetal CBD exposure does not affect latency to response to thermal stimulus in the Hargreaves test in wildtype or *TRPV1*^*KO/KO*^ female mice (A), even when controlling for estrus cycle (B). Fetal CBD exposure decreases latency to response in wild type CBD-exposed male mice (11.58 ± 0.64 seconds for vehicle-exposed vs 6.87 ± 3.27 seconds, P=4.993E-8, t-test), but does not affect latency response in *TRPV1*^*KO/KO*^ mice (11.089 ± 0.649 seconds vehicle-exposed, vs. 12.429 ± 1.610 seconds CBD-exposed P=0.453, t-test) (C). Graphs show fetal CBD exposure does not affect female offspring frequency in center zone (D), velocity (E), distance moved (F), or time in center zone, time moving, or time still (G) in the open field test. Graphs show that male offspring frequency in center zone (H), velocity (I), distance moved (J), or time in center zone, time moving, or time still (K) in the open field test. Graphs show fetal CBD exposure does not affect female offspring distance moved in open area (L), velocity (M), duration in open area, or zones (N), latency to enter the zones (O), or frequency entering zones (P), nor male offspring distance in open area (Q), mean velocity (R), duration in open area, or zones (S), latency to enter zones (T), or frequency entering zones (U) in light/dark box. The elevated zero maze shows that fetal CBD exposure does not affect female offspring velocity (V), frequency in light area (W), distance moved (X), time in closed or open areas (Y), frequency in closed area (Z), or zone crossings (AA), nor male offspring velocity (AB), frequency in light area (AC), distance moved (AD), time in closed or open areas (AE), frequency in closed area (AF), or zone crossings (AG).

### Fetal CBD exposure does not alter offspring anxiety

TRPV1 activity fetal development mediates offspring anxiety in mice^12,33^. Clinical studies show children exposed to whole cannabis in-utero have higher rates of anxiety and ADHD at puberty^15^. To determine if fetal CBD exposure affects offspring anxiety, we conducted the open field maze at six weeks old, the light dark box at eight weeks old, and the elevated zero maze test at ten weeks old (Figure 2D-AG). We found no differences in anxiety by any measure in male or female offspring based on fetal CBD exposure (statistics in supplemental table 1). Time in center zone of the open field test was not changed by fetal CBD exposure (90.20 ± 11.05 seconds, N=13 vehicle-exposed females versus 89.19 ± 6.36 seconds, N=25 CBD-exposed females, P = 0.933, t-test, 89.40 ± 10.05 seconds (N=19 vehicle-exposed males), 90.76 ± 6.00 seconds (N=23 CBD-exposed males), P=0.904, t-test. Duration in the open area of the light/dark box was equivalent between exposure groups (263.09 ± 17.52 seconds (N=6 vehicle-exposed females), 234.12 ± 11.89 seconds (N=21 CBD-exposed-females), P=0.225, Wilcoxon rank sum test. 233.82 ± 18.18 seconds (N=13 vehicle-exposed males), 246.27 ± 14.30 seconds (N=24 CBD-exposed males), P=0.239 Wilcoxon rank sum test. Time in the open area of the elevated zero maze test was equivalent between exposure groups (87.43 ± 9.16 seconds, (N=13 vehicle-exposed females), 93.41 ± 6.96 seconds, (N=25 CBD-exposed females), P=0.977, Wilcoxon rank sum test. 77.12 ± 6.22 seconds (N=23 vehicle-exposed males), 81.75 ± 5.76 seconds (N=24 CBD-exposed males), P=0.395, Wilcoxon rank sum test. Additional measures for each anxiety test can be found in Supplemental Table 1. We found significant differences between *TRPV1*^*KO/KO*^ and wild type mice, as previously characterized^33^ (Supplemental Figure 1). *TRPV1*^*KO/KO*^ mice also showed no differences in anxiety measures based on CBD exposure alone (Supplemental Figure 1). These data show that offspring anxiety as measured by the open field test, light/dark box, and elevated zero maze is not affected by fetal CBD exposure.

### Fetal CBD exposure does not affect offspring compulsivity

To determine the effect of fetal CBD exposure on offspring compulsivity, we conducted the Marble Burying Test. We found no significant differences in any measures of offspring compulsivity based on offspring sex or CBD exposure (46.14 ± 7.06 marbles buried N=14 vehicle-exposed females, 34.04 ± 3.63 marbles buried N=26 CBD-exposed females, P=0.098, t-test, 39.91 ± 4.40 marbles buried N=22 vehicle-exposed male, 42.09 ± 5.23 marbles buried N=23 CBD-exposed males, P=0.751, t-test). Additional measures from the marble burying test showed that fetal CBD exposure does not affect compulsivity (Supplemental Table 1).

### Fetal CBD exposure does not alter offspring spatial memory

To determine how fetal CBD exposure impacts offspring spatial memory, we conducted the Y maze test. We found no effect of CBD exposure, sex, or genotype (WT or *TRPV1*^*KO/KO*^) on spatial memory in the percent of correct alternations within the Y maze (Figure 3A/B, 0.656 ± 0.020 percent correct alternations N=13 vehicle females, 0.68 ± 0.025 percent correct alternations N=21 CBD females, P=0.63, t-test. 0.70 ± 0.021 percent correct alternations N=23 vehicle males, and 0.69 ± 0.027 percent correct alternations N=21 CBD males, P=0.699, t-test).

**Figure 3.**
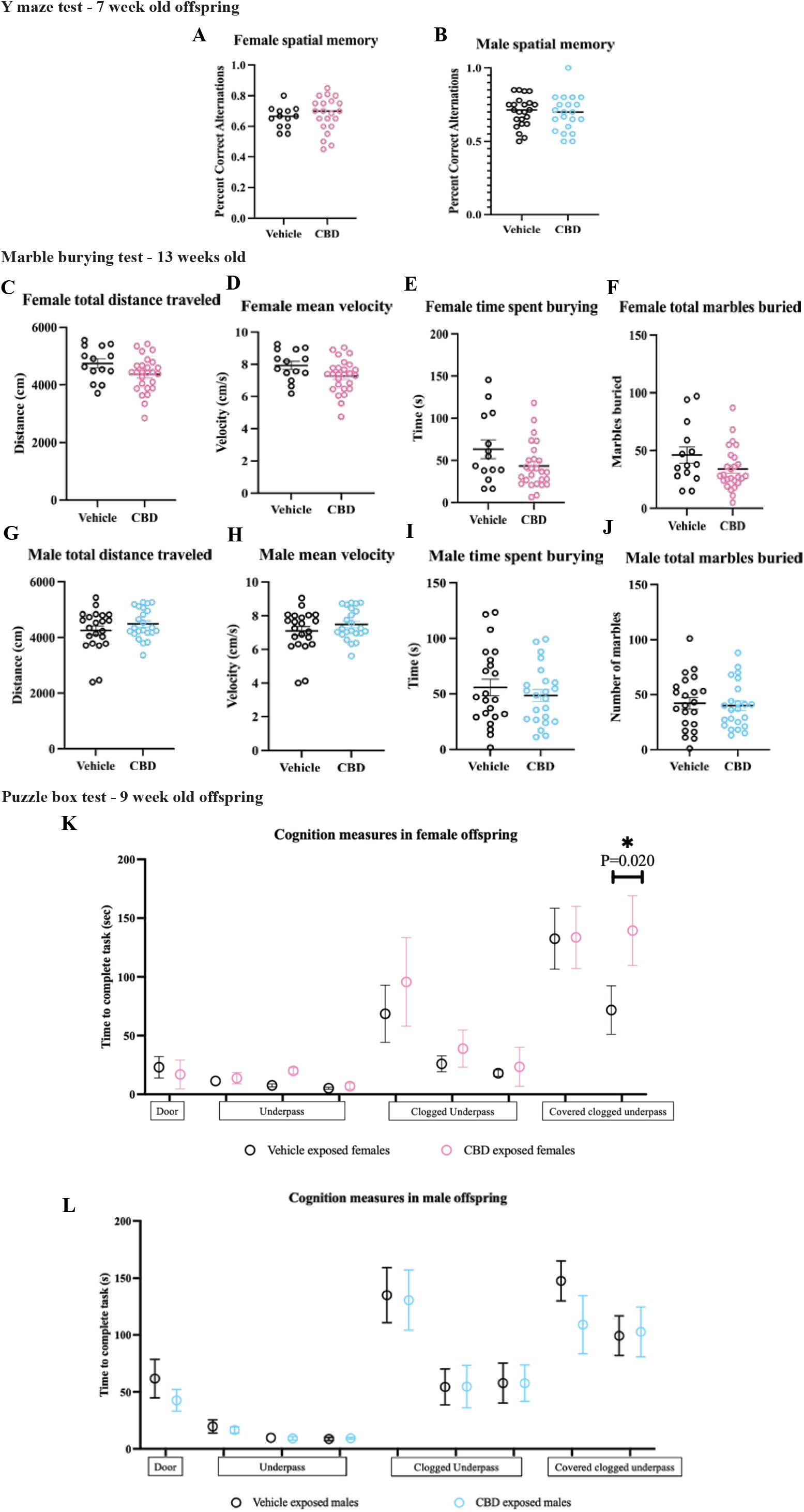
Fetal CBD exposure decreases female offspring cognition. Fetal CBD exposure does not affect female spatial memory (A) or male spatial memory (B) via the Y maze test. Graphs show fetal CBD exposure does not affect offspring compulsivity, including female total distance traveled (C), mean velocity (D), time spent burying (E) or marbles buried (F), nor male distance traveled (G), mean velocity (H), time spent burying (I) or total marbles buried (J), via the marble burying test. Graphs show fetal CBD exposure decreases female cognition at trial 9, (71.75 ± 20.71 seconds vehicle-exposed females, 139.42 ± 26.91 seconds CBD-exposed females, N=12 each, P=0.201, Wilcoxon rank sum test) (K), but not male cognition (L) via the puzzle box test.

### Fetal CBD exposure decreases cognition in female offspring

CBD can activate 5HT_1A_ receptors and Kv7 potassium channels. Fetal overactivation of serotonin receptors can negatively impact offspring cognition^20^. During fetal development, serotonin receptors are highly expressed in the prefrontal cortex (PFC), a region of the brain that mediates cognition^34^. Fetal overactivation of Kv7.2 and Kv7.3 is associated with intellectual disabilities, decreases in memory, and behavioral deficits^26^. To determine if fetal CBD exposure impacts offspring cognition, we conducted the puzzle box test. This test introduces each mouse to a light box and presents a progressively harder cognitive challenge to reach a dark goal area. Each mouse completes 9 trials, with novel progressive challenges at trials 2, 5, and 8. Mice with sufficient cognitive skills decrease their time to the goal area after secondary exposure to the obstacle. Male CBD-exposed offspring reached the goal box at all trials at similar times compared to vehicle-exposed male offspring (Figure 3L). Female CBD-exposed offspring took significantly more time to reach the goal area in trial 9 compared to the vehicle-exposed female offspring (71.75 ± 20.71 seconds vehicle-exposed females, 139.42 ± 26.91 seconds CBD-exposed females, N=12 each, P=0.02, Wilcoxon rank sum test), Figure 3K. These data show that fetal CBD exposure impairs cognition in female mice.

### Fetal CBD exposure decreases excitability of PFC L2/3 pyramidal neurons in a sex-dependent manner

We explored neural mechanisms that transduce fetal CBD exposure into decreased cognition in female mice (Fig. 3) with *ex vivo* electrophysiological recordings. We measured the intrinsic membrane properties of layer 2/3 pyramidal neurons in acute PFC slices from the CBD and vehicle-exposed male and female offspring. Fetal CBD-exposed female mice showed significantly decreased excitability (treatment effect, p<0.0001, two-way ANOVA), while CBD-exposed male pups were comparable to sex-matched vehicle treated controls (treatment effect, p=0.1711, two-way ANOVA; Fig. 4. 2A-C). Overall, fetal CBD exposure did not change the spike threshold (Vehicle: -39.84 ± 2.613 mV; CBD: -35.92 ± 1.523 mV; p=0.2018) (Fig. 4D). However, we observed a significant increase in both membrane potentials (Vehicle: 24.23 ± 2.251 mV; CBD: 33.21 ± 1.963 mV; p=0.0043) and minimum currents required to trigger action potentials (Vehicle: 110 ± 9.574 pA; CBD: 162.5 ± 11.36; p=0.0007) in CBD-exposed mice (Fig. 4D). We found that these differences stem from alterations in the intrinsic properties of female (Fig. 4F), but not male (Fig. 4H), offspring, without affecting resting membrane potentials (Vehicle: -64.07 ± 2.008 mV; CBD: -68.06 ± 1.773 mV; p=0.1261) (Fig. 4E, G, and I). Together, these data demonstrate CBD-mediated and sex-dependent decrease in neuronal excitability of PFC layer 2/3 pyramidal neurons.

**Figure 4.**
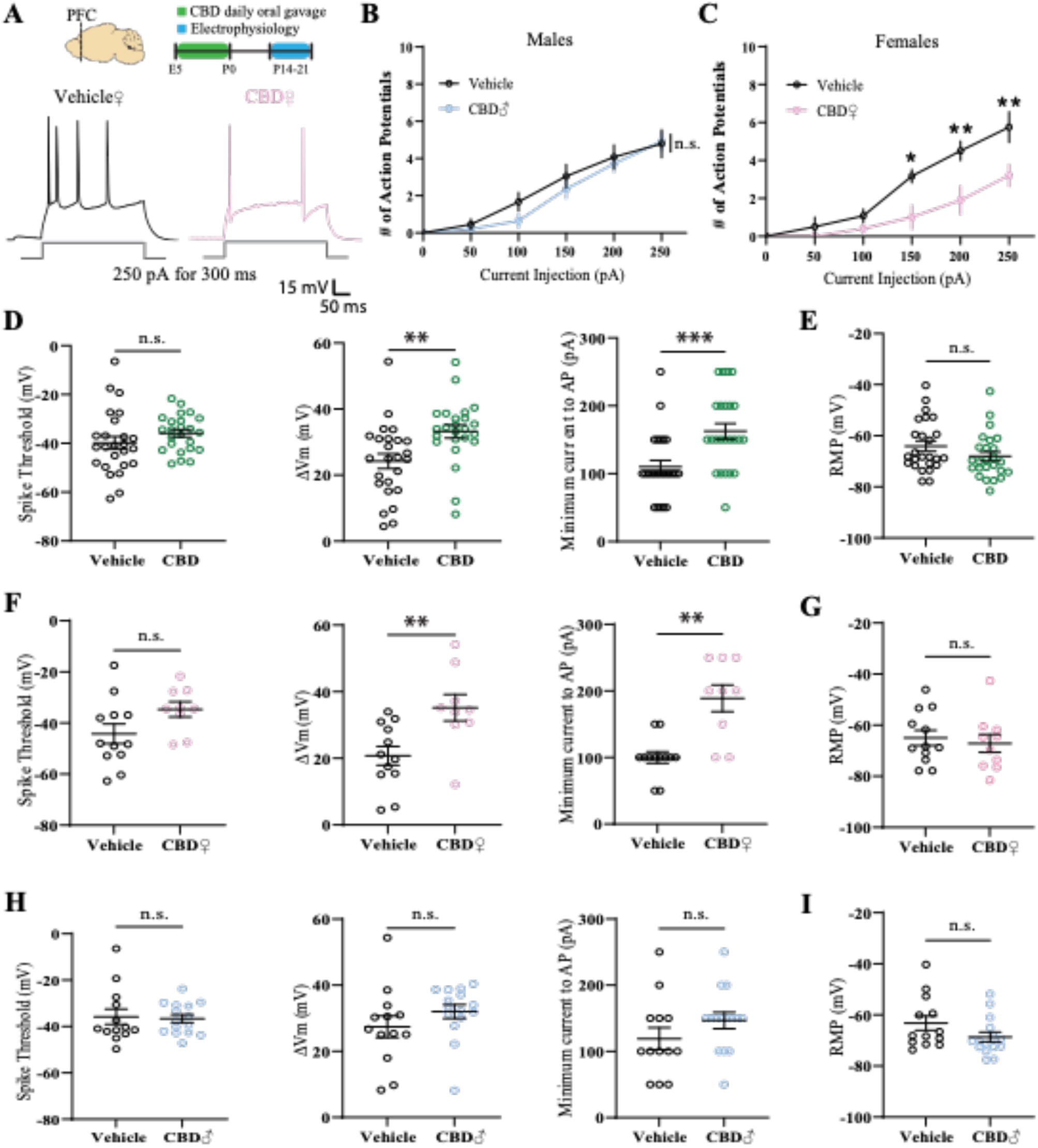
Fetal CBD exposure decreases excitability of PFC layer 2/3 pyramidal neurons in a sex-dependent manner. (A) Experimental timeline and representative traces of a stimulus train elicited by 250pA current injection for 300ms. (B and C) Fetal CBD exposure decreased the intrinsic excitability of layer 2/3 pyramidal neurons in P14 females but not males (vehicle females: n = 6 cells, 1 mouse; vehicle males: n = 12 cells, 2 mice; CBD females: n = 5 cells, 1 mouse; CBD males: n = 14 cells, 2 mice; female treatment effect: P < 0.0001; Sidak’s multiple comparison, * < 0.05, ** < 0.01). (D) Fetal CBD exposure did not alter spike thresholds of P14-21 mice (left, vehicle: -39.84 ± 2.613 mV, n = 25 cells, 4 mice; CBD: -35.92 ± 1.523 mV, n = 25 cells, 4 mice; Welch’s t-test, P = 0.2018). Fetal CBD exposure significantly increased membrane potential change (middle, vehicle: 24.23 ± 2.251 mV; CBD: 33.21 ± 1.963 mV; two-tailed t-test, P = 0.0043) and minimum currents (right, vehicle: 110 ± 9.574 pA; CBD: 162.5 ± 11.36; Mann Whitney, P = 0.0007) required to evoke action potentials. (E) Resting membrane potential of P14-21 mice remained unchanged following fetal CBD exposure (vehicle: -64.07 ± 2.008 mV, n = 25 cells, 4 mice; CBD: -68.06 ± 1.773 mV, n = 25 cells, 4 mice; Mann Whitney test, P = 0.1261). (F) The effect of fetal CBD exposure on changes of membrane potential (vehicle: 20.76 ± 2.826 mV, n = 12 cells, 2 mice; CBD: 35.19 ± 3.963, n = 10 cells, 2 mice; two-tailed t-test, P = 0.0066) and minimum current for action potential firing stemmed from females (vehicle: 100 ± 8.704 pA, n = 12 cells, 2 mice; CBD: 188.9 ± 20.03 pA, n = 10 cells, 2 mice; Welch’s t-test, P = 0.0018). (G) Resting membrane potential was unchanged in females following fetal CBD exposure (vehicle: -65.97 ± 2.966 mV, n = 12 cells, 2 mice; CBD: -67.16 ± 3.506 mV, n = 10 cells, 2 mice; two-tailed t-test, P = 0.6370). (H and I) Male mice showed no significant differences in spike threshold (vehicle: -35.81 ± 3.321 mV, n = 13 cells, 2 mice; CBD: -36.65 ± 1.725 pA, n = 15 cells, 2 mice; Mann Whitney test, 0.7856), membrane potential (vehicle: 27.43 ± 3.309 mV; CBD: 32.02 ± 2.115 mV; Mann Whitney test, P = 0.0648),and minimum currents for action potential spikes (vehicle: 119.2 ± 16.54 pA; CBD: 146.7 ± 12.41 pA; two-tailed t-test, P = 0.1893), or resting membrane potential (vehicle: -63.24 ± 2.820 mV; CBD: -68.67 ± 1.910, Mann Whitney test, P = 0.0977). *P < 0.05, **P < 0.01, ***P < 0.001; error bars represent SEM. n.s., not significant.

### Fetal CBD exposure affects excitatory synapse development in the PFC in a sex dependent manner

We next investigated whether fetal CBD exposure leads to sex-dependent structural and functional changes of excitatory spine synapses on layer 2/3 pyramidal neurons in the PFC because excitatory synapse development is regulated by neuronal activity^35,36,37^. We examined spine density (Fig. 5A and B) and function (Fig. 5A and D) in acute PFC slices of CBD and vehicle-exposed mice using two-photon microscopy and simultaneous whole-cell patch clamp recordings and two-photon glutamate uncaging^38^. We found that neither spine density (Vehicle: 0.91 ± 0.04 #/μm; CBD: 0.85 ± 0.04 #/μm; p=0.2513) nor size (Vehicle: 111.4 ± 8.07; CBD: 123.7 ± 6.76; p=0.2479) were affected by fetal CBD exposure (Fig. 5C). No sex-dependent changes were observed (Female, Vehicle: 0.90 ± 0.06 #/μm, CBD: 0.79 ± 0.07 #/μm, p=0.2813; Male, Vehicle: 0.906 ± 0.03 #/μm; CBD: 0.907 ± 0.02 #/μm; p=0.9750) (Fig. 5F and I). Surprisingly, however, uncaging-evoked alpha-amino 3-hydroxy-5-methyl-4 isoxazole propionic acid receptor (AMPAR) currents (uEPSCs) were significantly decreased on CBD-exposed groups (Fig. 5D and E) compared to vehicle treated controls (Vehicle: 7.49 ± 0.42 pA ; CBD: 6.09 ± 0.33 pA; p=0.011). We found this effect was female specific. uEPSCs were significantly smaller in fetal CBD-exposed female offspring (Vehicle: 8.66 ± 0.55 pA; CBD: 5.89 ± 0.51; p=0.0009) (Fig. 5G and H) with no effect on uEPSC amplitudes from CBD male mice (Vehicle: 6.48 ± 0.56 pA; CBD: 6.24 ± 0.45 pA; p=0.898) (Fig. 5J and K). Note that we targeted similar sizes of spines across groups as spine size and synaptic strength are strongly correlated^38^. These data show the female-specific effect of fetal CBD exposure on excitatory synapse development in the PFC.

**Figure 5.**
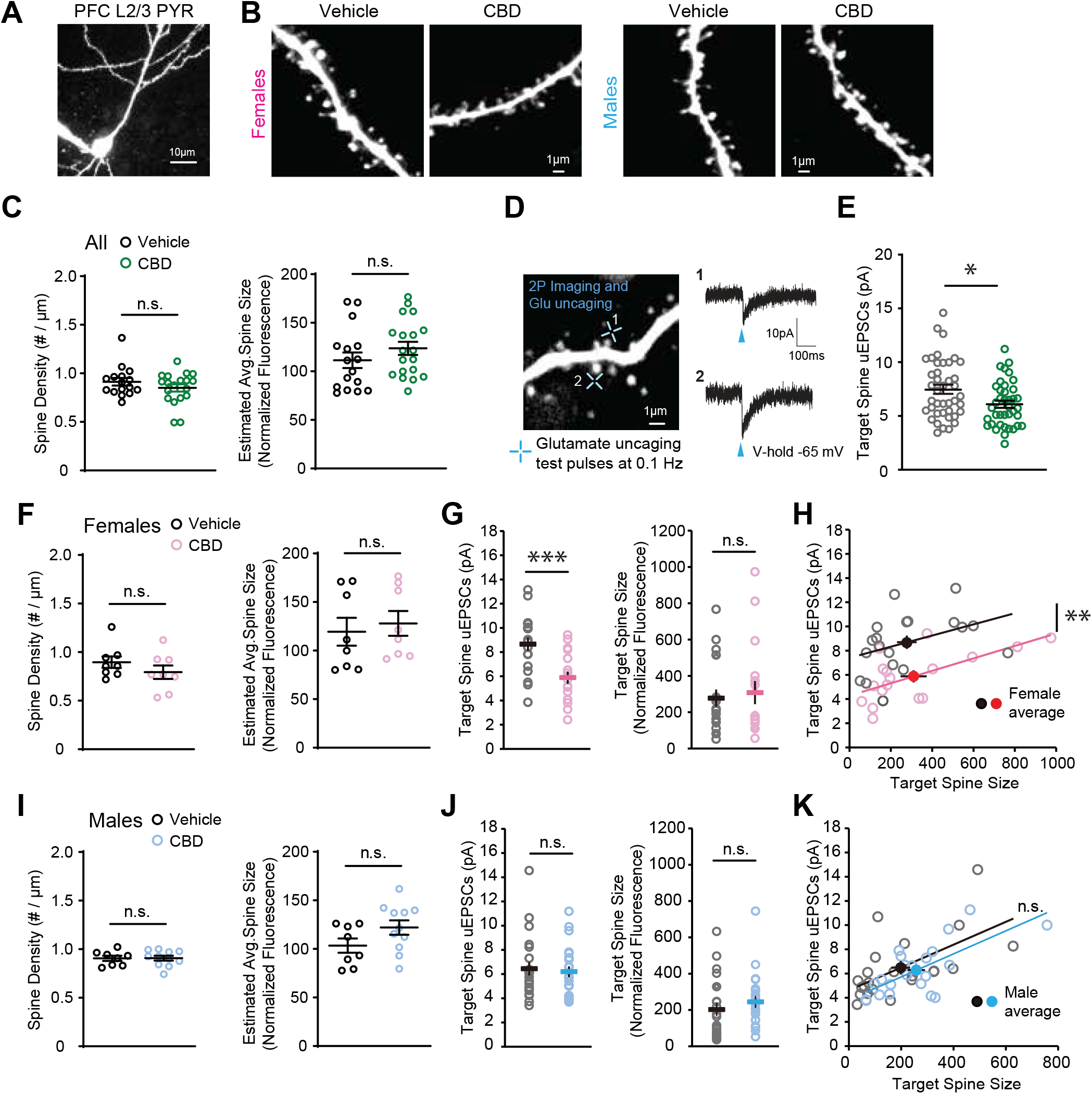
Fetal CBD exposure decreases synaptic strength of layer 2/3 pyramidal neurons in PFC of female mice. (A and B) Two-photon images of a whole-cell PFC layer 2/3 pyramidal neuron and dendritic segments from CBD and vehicle treated female and male mice at P14-22. (C) Fetal CBD exposure has no effect on spine density for combined male and female mice (vehicle: n = 70 dendrites, 16 cells, 4 mice; CBD n = 84 dendrites, 19 cells, 4 mice). (D) A two-photon image from a dendritic segment of PFC layer 2/3 pyramidal neuron and two-photon glutamate uncaging evoked EPSC (uEPSC) traces (average of 5 to 8 test pulses) recorded by whole-cell voltage-clamp recording (blue arrows indicate glutamate uncaging timepoint). (E) uEPSC amplitudes are significantly decreased in CBD-exposed offspring (vehicle: 7.49 ± 0.42 pA ; CBD: 6.09 ± 0.33 pA; P = 0.011, t-test) (vehicle: n = 41 spines, 14 cells, 3 mice; CBD n = 39 spines, 13 cells, 3 mice). (F) In female offspring, fetal CBD exposure has no effect on spine density or average spine size. (vehicle: n = 35 dendrites, 8 cells, 2 mice; CBD n = 36 dendrites, 8 cells, 2 mice). (G) uEPSCs recorded from similar sizes of target spines are significantly smaller in fetal CBD-exposed female offspring (vehicle: 8.66 ± 0.55 pA; CBD: 5.89 ± 0.51; P = 0.0009, t-test) (vehicle: n = 19 spines, 7 cells, 1 mouse; CBD n = 17 spines, 6 cells, 1 mouse). (H) Scatter plots showing significantly smaller uEPSCs in fetal CBD-exposed mice. (I) In male offspring, fetal CBD exposure had no effect on spine density or average spine size. (vehicle: n = 35 dendrites, 8 cells, 2 mice; CBD n = 48 dendrites, 11 cells, 2 mice). (J) In male offspring, CBD has no effect on uEPSCs (vehicle: n = 22 spines, 7 cells, 2 mice; CBD n = 22 spines, 7 cells, 2 mice). (K) Scatter plots showing comparable uEPSCs between fetal CBD-exposed and control mice. *P < 0.05, **P < 0.01; ***P < 0.001; error bars represent SEM. n.s., not significant.

## Discussion

CBD is easily accessible in many countries, and helps with nausea, the most common adverse symptom of pregnancy. The data presented here demonstrate that fetal CBD exposure sensitizes male offspring to thermal pain, decreases female offspring cognition, and reduces excitability of prefrontal cortical pyramidal neurons from female offspring. These results show CBD consumption during pregnancy can adversely affect fetal neurodevelopment. The data presented here are urgently needed to inform public health messaging.

### Fetal CBD exposure induces thermal pain sensitivity in male offspring

Our data demonstrate intrauterine CBD exposure increases thermal pain sensitivity in 11-week-old male offspring. CBD metabolites are not detected in pups after P8 suggesting that fetal CBD exposure alters thermal pain sensing circuits during development. This effect was dependent on the TRPV1 receptor. TRPV1 receptors are activated by high heat (40–45°C, 104-113°F)^39^ and are bound and activated by CBD^31^. TRPV1 overactivation by heat is hypothesized to be the mechanism that confers the detrimental fetal developmental effects of maternal fever, such as neural tube defects^11^. In a developmental context, altering the development of thermal pain circuits has critical postnatal implications. Persistent increased thermal pain sensitivity could increase the offspring’s susceptibility to chronic pain and could lay the ground for the use or dependency on pain relieving medications like opioids.

CBD-exposed female offspring responded to thermal stimuli similarly to vehicle-exposed controls. 17β-estradiol activation can downregulate TRPV1 activity in dorsal root ganglion sensory neurons^40^ and female mice show different thermal pain sensitivity across the estrus cycle^41^. Thus, it is possible that estrogen could protect female offspring from excessive activation of TRPV1 by intrauterine exposure to CBD.

Our data show that fetal CBD exposure increases thermal pain sensitivity in adult wild type male mice. Intrauterine CBD exposure reduced the latency to response in wild type male offspring compared to vehicle-exposed controls in the Hargreaves test. The Hargreaves test measures warm temperatures starting at 25°C/77°F with an increasingly hot light shined at the hind paw. CBD binds and activates TRPV1 and TRPV2^42^. While TRPV1 is activated by temperatures ranging from 40-45°C, 104-113°F, TRPV2 is activated by temperatures ranging from 50–53°C, 122-127.4°F^39^. *TRPV1*^*KO/KO*^ mice have similar thermal sensitivity to wild type mice when measured using the Hargreaves test here and in previously published studies, but thermal sensitivity of *TRPV1*^*KO/KO*^ mice can be distinguished from wild type with the 50°C/122°F hot plate^43^. Intrauterine CBD exposure does not impact thermal sensitivity in *TRPV1*^*KO/KO*^ mice, suggesting that the effect of CBD on thermal pain circuit development depends on TRPV1. These results suggest that excessive activation of TRPV1 during fetal development can alter long term thermal sensitivity.

### Fetal CBD exposure does not impact offspring anxiety or compulsivity

The prefrontal cortex is a region of the brain that controls cognition, memory, anxiety, attention, and impulsivity^44^. The developing prefrontal cortex contains a multitude of receptors critical for normal development, including 5HT_1A_ serotonin receptors and Kv7 receptors that are bound and activated by CBD^45,46^. We investigated multiple behaviors mediated by the prefrontal cortex, including anxiety and compulsivity. CBD-activated receptors, including TRPV1, are expressed in the hippocampus, a region of the brain that mediates memory^13^.

We found that intrauterine CBD exposure did not impact offspring anxiety in wild type or *TRPV1*^*KO/KO*^ mice by any measure in the open field test, the light dark box, or the elevated zero maze test. Previous studies administered 20mg/kg CBD (Epidiolex) dissolved in honey via oral gavage from 14 days pre-conception through offspring weaning and found that the 12-week-old CBD-exposed female offspring buried more marbles in the marble burying test while male CBD-exposed offspring were no different than control^47^. In contrast, we found that oral gavage of 50mg/kg CBD from E5 through birth did not significantly affect anxiety measured by the open field, light/dark box, or elevated zero maze, nor any measures of compulsivity on the marble burying test.

### Fetal CBD exposure decreases cognition in female offspring, but not male offspring

We show fetal CBD exposure reduces cognition in female offspring. Cognition is mediated by the prefrontal cortex^44^. We show that fetal CBD exposure reduces excitability of P14-P21 pyramidal neurons from the female prefrontal cortex. Fetal CBD exposure raised the required current to elicit an action potential, raised the required mV to elicit an action potential, and decreased the number of action potentials elicited at set current.

The electrophysiological and cognitive effects of fetal CBD exposure could be mediated by excessive activation of two different ion channel receptors. CBD activates 5HT_1A_ and Kv7.2/3, both of which mediate neuronal activity^8,10^ and are expressed in the fetal prefrontal cortex^45,29^ (Figure 6A). When 5HT_1A_ is excessively activated during development, offspring show decreased neurogenesis, decreased neuronal activity in corticotropin releasing neurons, decreased neuron network complexity, changes in neuronal refinement, decreases in the amplitude of sensory-evoked potentials, a delay in sensory evoked potentials, and decreased sensory and spontaneously evoked firing^20^. There is evidence that Kv7 channels also regulate neuronal activity. For example, Kv7.2/3 gain of function mutations are associated with epilepsy and intellectual disability in humans^26^. Excessive activation of Kv7.2/3 decreases the refractory period following action potentials and increases post-conditioned super excitability of neurons^48,49^. Because fetal CBD exposure impacts neuronal excitability and cognition and activates 5HT_1A_ and Kv7.2/3, which mediate neuronal development and excitability, excessive activation of 5HT_1A_ and Kv72/3 could be a mechanism by which CBD alters cognition. In contrast, fetal CBD exposure does not alter resting membrane potential, which is not set by 5HT_1A_ nor Kv7.2/3. The extent to which excessive activation of 5HT_1A_ and Kv7.2/3 contribute to cognitive and cellular effects of intrauterine CBD exposure on female mice will be an avenue of future studies.

**Figure 6.**
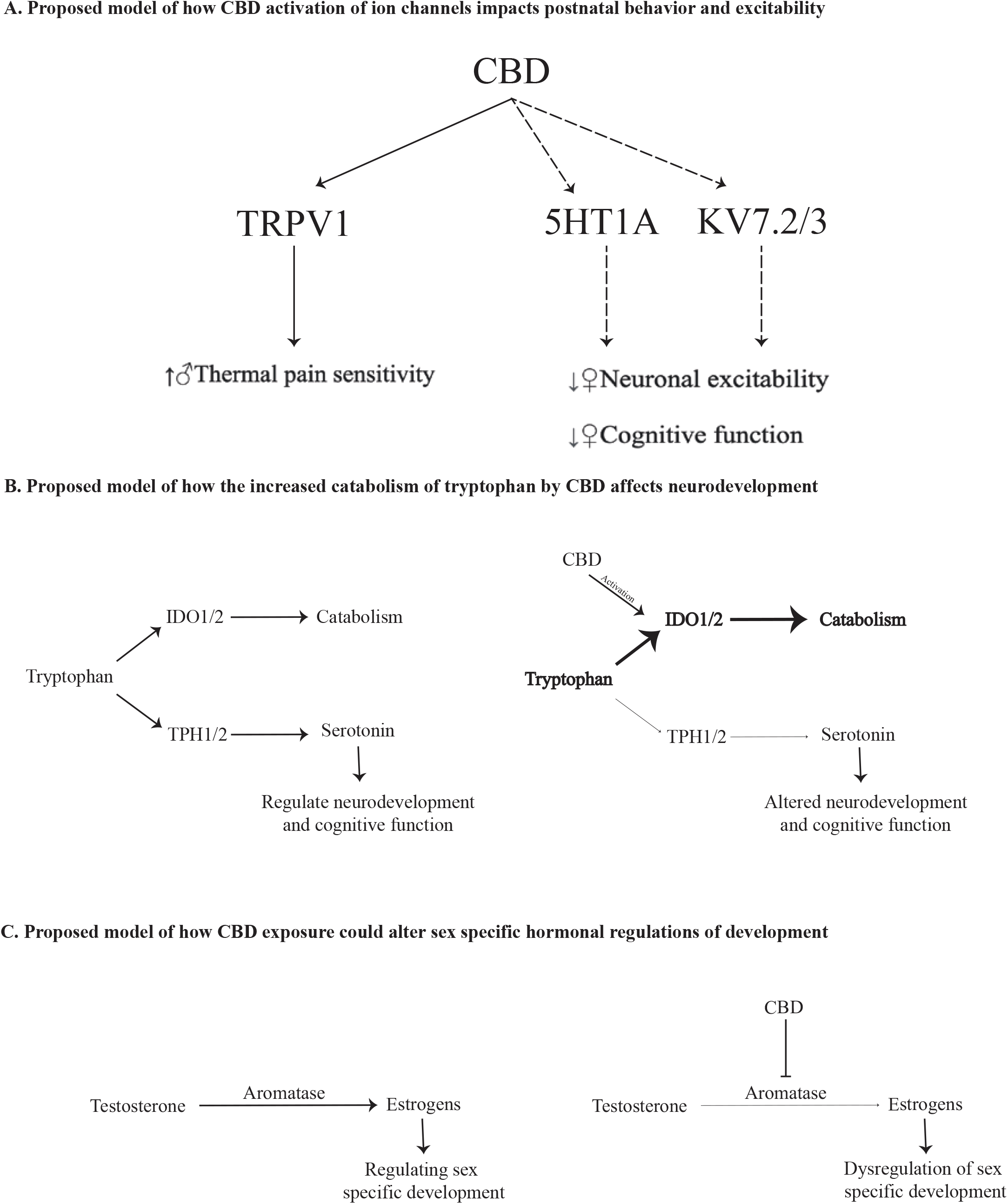
Model of effects of fetal CBD exposure. Fetal CBD exposure increases male thermal pain sensitivity, decreases female cognition, and alters female prefrontal cortex pyramidal neurons. CBD activates TRPV1, 5HT_1A_, and Kv7.2/3 receptors which each modulate effects of CBD that we observed (A). CBD activates the IDO1/2 pathway which could increase catabolism of tryptophan thereby reducing levels of serotonin to alter cognition (B). CBD inhibits aromatase which would reduce estrogen levels to potentially affect sex specific development.

CBD may alter female cognition through its effect on serotonin production (Figure 6B). CBD activates catabolism of tryptophan, a precursor of serotonin, which could deplete endogenous serotonin. Reduction of serotonin levels can reduce cognitive function and alter neuronal excitability in mice^50,51^. Tryptophan depletion impairs memory consolidation in humans^21^ and hinders cognitive function in patients with Alzheimer disease^52^. In rats, tryptophan depletion impairs object recognition^53^. Genetic deletion of tryptophan hydroxylase 2 (TPH2), the enzyme that converts tryptophan to serotonin, impairs learning and cognitive flexibility in mice^22^.

The sex specific effects of CBD on neuronal excitability and cognition may be due to one or multiple intertwined pathways. Female mice have higher levels of circulating 5HT than do males, and higher expression of 5HT_1A_^54,55,56^. CBD may reduce circulating levels of estrogen because CBD inhibits aromatase, the enzyme that converts androgens to estrogens^57^ (Figure 6C). Aromatase inhibition increases male cognition in mice^58^ and humans with aromatase polymorphisms have increased risks of age-related cognitive decline^59^. In clinical studies, reduction of estrogen levels is associated with decreased female cognition, learning, and memory^60^. Decreases in circulating estrogens also mediates offspring neurodevelopment and behavior^61^. Reduction of estrogen levels decreases neuronal survival, differentiation, and plasticity, and downregulates the cholinergic and glutamate systems in mice and nonhuman primates^62,60^. Thus, one potential mechanism by which fetal CBD exposure elicits its effects on cognition could be through reduction of circulating estrogens which mediate cognition in female mice. Mechanisms by which intrauterine CBD exposure impacts cognition and thermal sensitivity in a sex specific manner will be exciting new avenues for future studies.

Cannabis consumption during pregnancy is increasing^2^. Pregnant patients can self-medicate nausea symptoms with whole cannabis or CBD alone. Clinical studies show fetal cannabis exposure is associated with adverse behavioral outcomes^63^. Most cannabis products contain CBD. There is likely an additional population of pregnant people who consume CBD alone because it is not psychoactive. There is a gap in understanding the effect of gestational CBD consumption, despite high consumption rates. Our work shows that a high dose fetal CBD exposure increases male thermal pain sensitivity, reduces excitability of pyramidal neurons in the prefrontal cortex in female mice, and decreases female cognition. This research is urgently needed to inform public health messaging that CBD consumption during pregnancy can have adverse long-term neurodevelopmental outcomes.

The data presented here show fetal CBD exposure poses risks to offspring neurodevelopment and behavior. CBD exposure increased thermal pain sensitivity for male offspring and reduced cognition in female offspring. CBD exposure reduced excitability of the prefrontal cortex. We show that CBD exposure alone cannot account for some of the human behavioral outcomes associated with gestational cannabis exposure including anxiety, compulsivity, and spatial memory. This data fills a critical gap in the translational research focused on gestational cannabis consumption. Clinical work cannot distinguish between cannabis component parts. As CBD gains traction as a remedy for morning sickness, it is critical that we reveal the effects of gestational CBD exposure. This work should be used in the clinical sphere to caution pregnant people against CBD consumption during pregnancy. Further research to determine critical periods of CBD exposure and mechanisms of action are urgently needed to inform public health messaging and clinical practice.

## Methods

### Study design

C57Bl6J female mice were administered 50mg/kg of CBD (NIDA) dissolved in sunflower oil or sunflower oil alone by oral gavage from E5 though birth. Researchers were blinded through the entirety of behavioral testing as to which group was CBD-exposed and which was control. At 21 days old, offspring were weaned into standard chow, and cohoused with their same-sex siblings. Our sample size includes 27 vehicle-exposed and 27 CBD-exposed dams, whose litter sizes vary (Figure 1). All experiments were approved by the University of Colorado Anschutz Medical Campus Institutional Animal Care and Use Committee. Each behavior experiment was performed once per animal, within a two-week period to accommodate high volumes of offspring. Both exposure groups were represented in each individual trial. The objectives of our study are to understand the effect of fetal CBD exposure on offspring thermal pain sensitivity, anxiety, spatial memory, compulsivity, cognition, and prefrontal cortex excitability.

### Animal protocols

These experiments were approved by the University of Colorado Anschutz Medical Campus Institutional Animal Care and Use Committee (protocol #139). Female C57BL6 mice (Strain #000664) and female *TRPV1*^*KO/KO*^ mice (Strain #003770) on a C57BL6 background (Jackson Laboratory, Maine) Females were individually housed mated with a single male. Upon visualization of a vaginal plug (E0.5), the male mouse was removed. Dam weight was tracked each day starting on E0.5. Any mouse that had not gained appropriate weight by E14 was removed from the study.

### CBD administration

Cannabidiol (CBD) (98.7% pure powder, synthetic, National Institutes of Drugs of Abuse) was obtained following approval of our Drug Enforcement Administration (DEA) Schedule 1 Drug license. 500mg of CBD was diluted in 40 ml sunflower oil and heated to 60°C to make a 12.5 mg/ml concentration in an amber glass vial to avoid light exposure. A second identical vial was filled with equal volume of sunflower oil. Vials were labeled drug “A” and “B” by author E.A. Bates to allow author K.S. Swenson to remain blinded for the duration of behavior experiments. Diluted CBD was evaluated for purity by the iC42 lab at the University of Colorado Anschutz Medical Campus. Consumption method (injection, oral consumption, inhalation) affects pharmacokinetic CBD breakdown. CBD is most commonly consumed topically (lotions, balms) or orally^30^. Oral consumption of CBD has a slower pharmacodynamic clearance than with IP injection^64^. CPY3A4 metabolizes CBD into the primary active metabolite 7-OH-CBD. CBD is metabolized by CYP3A4 to the primary active metabolite of 7-OH-CBD. This first pass metabolism reduces the bioavailability of CBD to 10-13% of the initial dose^65^. We chose to multiply the standard research dose (5mg/kg administered as i.p. injection) by 10 to create a comparable oral dose of 50mg CBD/kg body weight. This dose is well below the dose that would induce hepatotoxicity when administered via oral gavage repeatedly over multiple days^66^. CBD is highly lipophilic and easily crosses the placenta into the fetal blood stream, where it accumulates in fat-heavy organs such as the brain and liver^6^.

Data collected on the dams included daily weighing, gestational length, litter size, and pup vitality. We calculated an estimated average pup weight (last pregnant day weight – pre-pregnancy weight) / litter size) for each dam. All data points were analyzed for sex differences whenever possible. Unblinding to exposure group occurred after all analyses were completed.

### Plasma metabolite concentration

On E18.5, dams were euthanized via isoflurane inhalation and secondary cervical dislocation. The uterus was removed, opened, and pups were separated from the uterus. Pups were removed from their placentas, decapitated, and blood was collected into EDTA tubes. P4, P8, and P12 pups were euthanized, and blood collected into EDTA tubes. Dam blood was collected via decapitation or cardiac puncture and blood was stored in EDTA tubes (Microvette 100 KE3 Kent Scientific Corporation, item ID: MCVT100-EDTA). Blood was stored on ice and centrifuged at 4°C at 3000xg for 10 minutes to separate plasma. Plasma was stored in a clean EDTA tube at −80°C until transfer to the iC42 Clinical Research and Development (Aurora, CO). CBD, 6a-hydroxy-CBD, 7-hydroxy-CBD, carboxy-CBD, and CBD glucuronide were quantified using high-performance liquid chromatography-tandem mass spectrometry (LC-MS/MS) as previously described^67^. The results included in the study sample batch met predefined acceptance criteria: the calibration range for 6a-hydroxy-CBD, 7-hydroxy-CBD and carboxy-CBD are 1.56-400 ng/mL, CBD range was 0.39-400ng/ml, and CBD-glucuronide range was 0.78-200ng/ml. There was no carryover and no matrix interferences. Accuracy in the study sample batch was within the ±15% acceptance criterion and imprecision was <15%.

### Behavior

All behavior protocols were obtained from the University of Colorado Anschutz Medical Campus Animal Behavior Core staff. These tests occurred in controlled light, temperature, humidity, silent, pathogen-free environment. Unless otherwise noted, behavioral tests were completed between 9AM and 1PM. All mice were tested following schedule: open field test at 6 weeks of age, y maze test at 7 weeks of age, light dark box test at 8 weeks of age, puzzle box test for cognition at 9 weeks of age, elevated zero maze test for anxiety at 10 weeks of age, marble burying for compulsivity at 13 weeks of age, and Hargreaves for thermal pain sensitivity at 11 weeks of age. Open field, light/dark box, elevated zero maze, and puzzle box data were collected and analyzed using the Ethovision XT software from Noldus using version 8.5. Before and after every mouse, any behavior equipment was wiped to remove soiling followed by 70% ethanol wiping. In any experiments involving multiple chambers which run multiple mice at a time, individual chambers were used sequentially for every mouse from one cage before being replaced with sex-matched mice from another cage to reduce any stress due to lingering scents.

### Hargreaves test

The Hargreaves test is a commonly used method to test murine temperature sensitivity. We placed each mouse in a glass bottom enclosure heated to 30°C/86°F temperature to minimize errors arising from heat sink effects. Mice were habituated to the apparatus for 1 hour the day before the test, and at least 30 minutes before any testing began. Once the mouse was standing still in the enclosure, we applied an infrared heat source under the plantar surface of the hind paw and quantified the latency to response to a heat source. The intensity of the heat source was set to 10 amps which produced withdraw latencies of 5-15 seconds in naïve animals, which allows enough time for heat detection and response but is short enough to not cause thermal damage. We tested eighteen CBD-exposed and twenty-one vehicle exposed female mice from seven different litters per exposure and eight CBD-exposed and nine vehicle exposed male mice from three different litters per exposure.

### Estrus Cycle Tracking

The vaginal cytology method was used to track the estrus cycle of female mice^68^. Immediately after the Hargreaves test, female mice were lavaged to collect cells from the vaginal wall. PBS was pipetted up and down 3 times within the vagina to obtain cells. Cells were mounted on a dry slide, overlaid with a coverslip, and immediately viewed at 200X magnification under bright field illumination. Estrus stage was determined based on the presence or absence of leukocytes, cornified epithelial, and nucleated epithelial cells^69^.

### Open Field Test

The open field test measures murine anxiety and locomotor activity. We place the mice in a 44Wx44Lx25H cm arena under bright lighting conditions (900-1000lux) for 10-minute sessions each. Ethovision tracking system measures total distance traveled in centimeters along with time spent in the outer and central zones of the box. We tested 14 vehicle-exposed female offspring from 5 litters, 25 CBD-exposed female offspring from 7 litters, 18 vehicle-exposed males from 6 litters, and 21 CBD-exposed males from 7 litters in the Open Field Test.

### Light Dark Box

The light dark box tests murine unconditioned anxiety. The mice were placed in a box (45WX22.5LX28H cm) with one dark, covered section and one lit, open section, separated with a dark wall containing a door. Each mouse was placed into the closed section of the box for 5 minutes, then the door was removed, and the mouse was able to explore the open area under video monitoring for 5 minutes. We quantified time spent in the open and closed areas, and number of transitions between the two areas. We tested six vehicle-exposed female offspring from three litters, nineteen CBD-exposed females from five litters, thirteen vehicle-exposed male offspring from four litters, and twenty-two CBD-exposed male offspring from six litters in the light/dark box.

### Elevated Zero Maze

The elevated zero maze tests murine anxiety. Mice were placed individually on a circular runway (50cm diameter, 5cm wide track, 50cm above ground) which is divided into four 90° quadrants. Two opposing quadrants are surrounded by 30cm high walls while the in-between quadrants have no walls. The mice were placed facing the entrance of one of the walled quadrants. Time spent in each quadrant was video recorded and scored for the number of zone transitions, distance moved, and percentage of time in open and closed zones. Each mouse is tested for ten minutes. We tested thirteen vehicle-exposed female offspring from five litters, twenty-five CBD-exposed female offspring from seven litters, twenty-three vehicle-exposed male offspring from seven litters, and twenty-six CBD-exposed male offspring from seven litters in the elevated zero maze.

### Y Maze Test

The y maze spontaneous alternation test quantifies murine spatial cognition. Mice were placed in the center of a y-shaped maze with three opaque arms at 120° angles from each other. The mouse was free to explore all three arms of the maze. Mice were monitored for either 10 minutes or 22 arm-changes, or whichever happened first. We calculate the percentage of “correct” and “incorrect” movements, where correct patterns are three subsequent arm changes (e.g., arm A to arm B to arm C) and incorrect patterns are three arm changes in repeated arms (e.g., arm A to arm B to arm A). Entry to the arm is marked once all four limbs have entered that arm. We tested thirteen vehicle-exposed female offspring from five litters, twenty-one CBD-exposed female offspring from seven litters, twenty-three vehicle-exposed male offspring from seven litters, and twenty-one CBD-exposed male offspring from seven litters in the Y Maze.

### Marble Burying Test

Marble burying measures compulsivity in mice. Our apparatus is an 11cmX11cm box filled with a layer of bedding and a 3×3 square of evenly placed blue marbles on top of the bedding under recording on the Ethovision video monitoring system. Mice were placed in the apparatus for 10 minutes. We quantified the total distance traveled and velocity of the mice from the Ethovision tracking system, and marble burying was quantified manually. We quantified the number of marbles buried, number of marbles re-buried, and time spent burying (seconds and percentage of total time). Apparatuses were reset between each mouse from the same cage, through between mice from separate cages we removed the marbles and bedding, cleaned the boxes and marbles with 70% ethanol, and replaced the setup with fresh bedding. When possible, littermate sex-matched mice were tested in the same box to reduce stress caused by any latent smells. We tested fourteen vehicle-exposed female offspring from five litters, twenty-six CBD-exposed female offspring from seven litters, twenty-two vehicle-exposed male offspring from seven litters, and twenty-three CBD-exposed male offspring from seven litters.

### Puzzle box

The puzzle box measures murine cognition. The puzzle box is a Plexiglas white box divided by a removable barrier into two compartments: a brightly lit start zone (58 cm long, 28 cm wide) and a smaller covered goal zone (15 cm long, 28 cm wide). Mice are motivated to move into the goal zone by their aversion to the bright light in the start zone. We placed individual mice into the start zone and measured the time to move through the 4cm wide underpass to the goal zone (dark compartment). Each mouse underwent nine trials (T1-T9) over the course of three days, with three trials each day. Each day the underpass was obstructed with increasing difficulty. For T1 (training) the underpass is clear, and the barrier has an open door over the location. On T2 and T3 (day 1) and T4 (day 2), the mice go through an underpass. On T5 and T6 (day 2) and T7 (day 3), the mice must dig through the sawdust-filled underpass to reach the goal zone. During T8 and T9 (day 3), the mice must remove a 4×4cm covering and then dig through sawdust to reach the goal zone. This sequence allows assessment of problem-solving abilities (T2, T5 and T8), learning/short-term memory (T3, T6, and T9), and repetition on the next day provides a measure of long-term memory (T4 and T7). We tested twelve vehicle-exposed female offspring from five litters, twelve CBD-exposed female offspring from five litters, twelve vehicle-exposed male offspring from four litters, and twelve CBD-exposed male offspring from four litters in the puzzle box.

### Preparation of acute prefrontal cortex (PFC) slices

Acute coronal PFC slices were obtained from P14 to 22 C57BL/6 male and female wild-type mice prenatally exposed to either CBD or vehicle in accordance with the Institutional Animal Care and Use Committees of the University of Colorado on Anschutz Medical Campus and National Institutes of Health guidelines. Mice were anesthetized with isoflurane and euthanized by decapitation. Immediately after decapitation, the brain was extracted and placed in icy cutting solution containing 215mM sucrose, 20mM glucose, 26mM NaHCO_3_, 4mM MgCl_2_, 4mM MgSO_4_, 1.6mM NaH_2_PO_4_, 1mM CaCl_2_, and 2.5mM KCl. Using a Leica VT1000S vibratome, the PFC was sectioned into 300μm thick slices. PFC slices were incubated at 32°C for 30 minutes in 50% cutting solution and 50% artificial cerebrospinal fluid (ACSF) composed of 124mM NaCl, 26mM NaHCO_3_, 10mM glucose, 2.5mM KCl, 1mM NaH_2_PO_4_, 2.5mM CaCl_2_, and 1.3mM MgSO_4_. After 30 minutes, this solution was replaced with ACSF at room temperature. For all two-photon and electrophysiology experiments, the slices were placed in a recording chamber and bathed in carbogenated (95% O_2_ / 5% CO_2_) ACSF at 30°C and allowed to equilibrate for at least 30 minutes prior to the start of experiments.

### Two-photon Imaging

Two-photon imaging was performed on layer 2/3 pyramidal neurons at depths of 20-50 μm of PFC slices at P14-P22 using a two-photon microscope (Bruker) with a pulsed Ti:sapphire laser (MaiTai HP, Spectra Physics) tuned to 920 nm (4-5 mW at the sample). All experiments were controlled using the Praire View (Bruker) software. Neurons were imaged at 30°C in recirculating ACSF with 2mM CaCl_2_, 1mM MgCl_2_ aerated with 95% O_2_ / 5% CO_2_. For visualization, cells were whole-cell patched and filled with Alexa 488. For each neuron, image stacks (512 × 512 pixel; 0.047 um/pixel) with 1-μm z-steps were collected from secondary or tertiary distal apical dendrites. All images shown are maximal projections of three-dimensional image stacks after applying a median filter (2 × 2) to the raw image data. All protrusions on the dendritic shaft were counted as dendritic spines in images of green (Alexa 488) channel using ImageJ software (NIH). Dendritic spines density was calculated by dividing the number of spines by the dendritic length (in μm). Spine size was estimated from background-subtracted and bleed-through-corrected integrated pixel fluorescence intensity of the region of interest (∼1μm^2^) surrounding the spine head. This measurement was normalized to the mean fluorescence intensity of the dendritic segment adjacent to the dendritic spine of interest^35,70^.

### Electrophysiology and Two-photon glutamate uncaging

PFC layer 2/3 pyramidal neurons were identified by morphology and distance from slice surface (< 40μm), and patched in the whole-cell (4-8 MΩ electrode resistance; 20-40 MΩ series resistance) current clamp configuration (MultiClamp 700B, Molecular Devices). Using a potassium-based internal solution (136mM K-gluconate, 10mM HEPES, 17.5mM KCl, 9mM NaCl, 1mM MgCl_2_, 4mM Na_2_-ATP, 0.4mM Na-GTP, 0.2mM Alexa 488, and ∼300 mOsm, ∼pH 7.26), spiking properties of layer 2/3 neurons were examined at 30°C in recirculating ACSF with 2mM CaCl_2_, 1mM MgCl_2_. Excitability was measured by injection of depolarizing current steps (50 - 250pA, 300ms). Resting membrane potential was recorded prior to the first depolarizing current step. Minimum current required to elicit an action potential was defined as the smallest current step that triggered at least one spike. The spike threshold was defined as the potential at which the spike is triggered. The change in potential (ΔVm) was defined as the difference (absolute value) between the resting membrane potential and spike threshold. For two-photon glutamate uncaging experiments, whole-cell recordings in the voltage clamp configuration (V_hold_ = - 65mV) were performed in 1μM TTX and 2.5mM MNI-glutamate (Tocris) containing ACSF at 30°C. Using cesium-based internal solution (135mM Cs-methanesulfonate, 10mM HEPES, 10mM Na_2_ phosphocreatine, 4mM MgCl_2_, 4mM Na_2_-ATP, 0.4mM Na-GTP, 3mM Na L-ascorbate, 0.2mM Alexa 488, and ∼300 mOsm, ∼pH 7.25), individual dendritic spines (on secondary or tertiary apical dendritic branches, 50-100μm from soma) were targeted by glutamate uncaging and uncaging evoked excitatory postsynaptic currents (uEPSCs) were recorded^35^. uEPSC amplitudes were quantified as the average (5-10 test pulses at 0.1 Hz) from a 1 ms window centered on the maximum current amplitude within 30 ms after uncaging pulse delivery.

### Statistical Analyses

Unless otherwise specified, all data were collected and segregated by sex, exposure group, and genotype. We first completed a D’Agostino and Pearson test to determine if data were normally or nonnormally distributed. When normally distributed, data within each direct comparison (two exposure groups within one sex and one genotype) were analyzed via a t test, and any with a p-value less than 0.05 is reported as significant. If data were nonnormally distributed, we completed a Wilcoxon rank sum test for data within each direct comparison and reported any significant data with a p value less than 0.05.

---

**Supplementary table 1.**
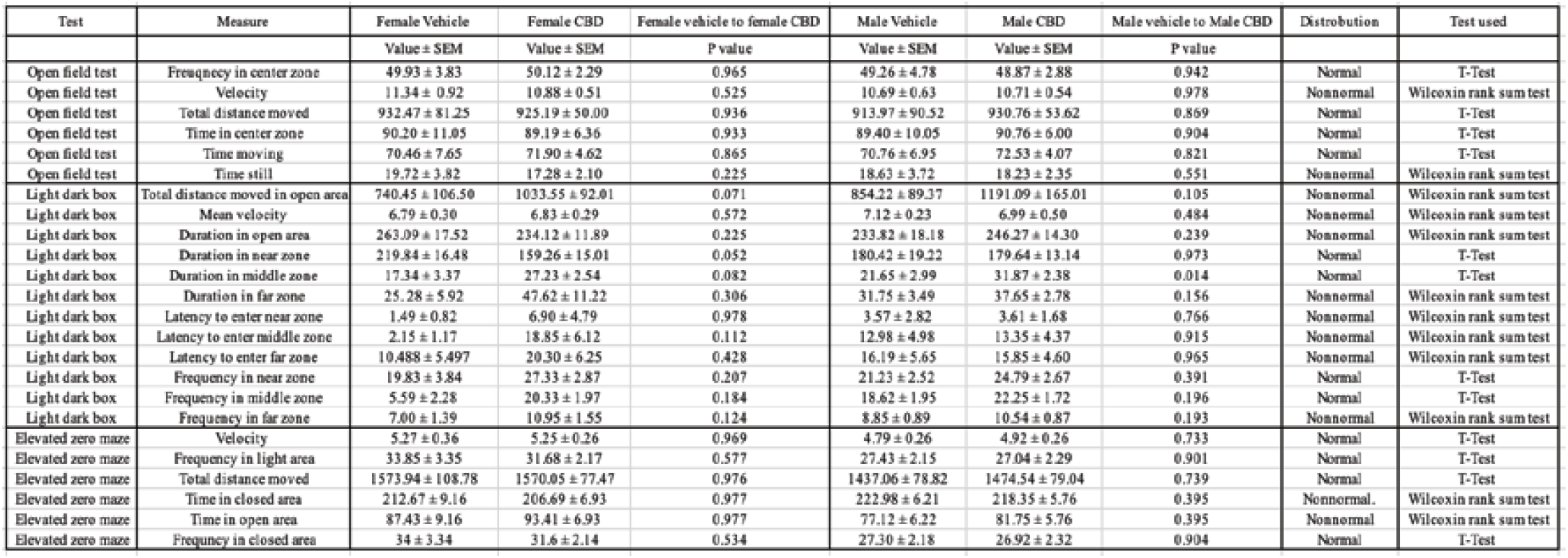
Data for anxiety tests. **Summary of anxiety measure statistics.** Table shows all measures for the open field test, light dark box, and elevated zero maze split by sex and statistical analysis.

**Supplemental Figure 1.**
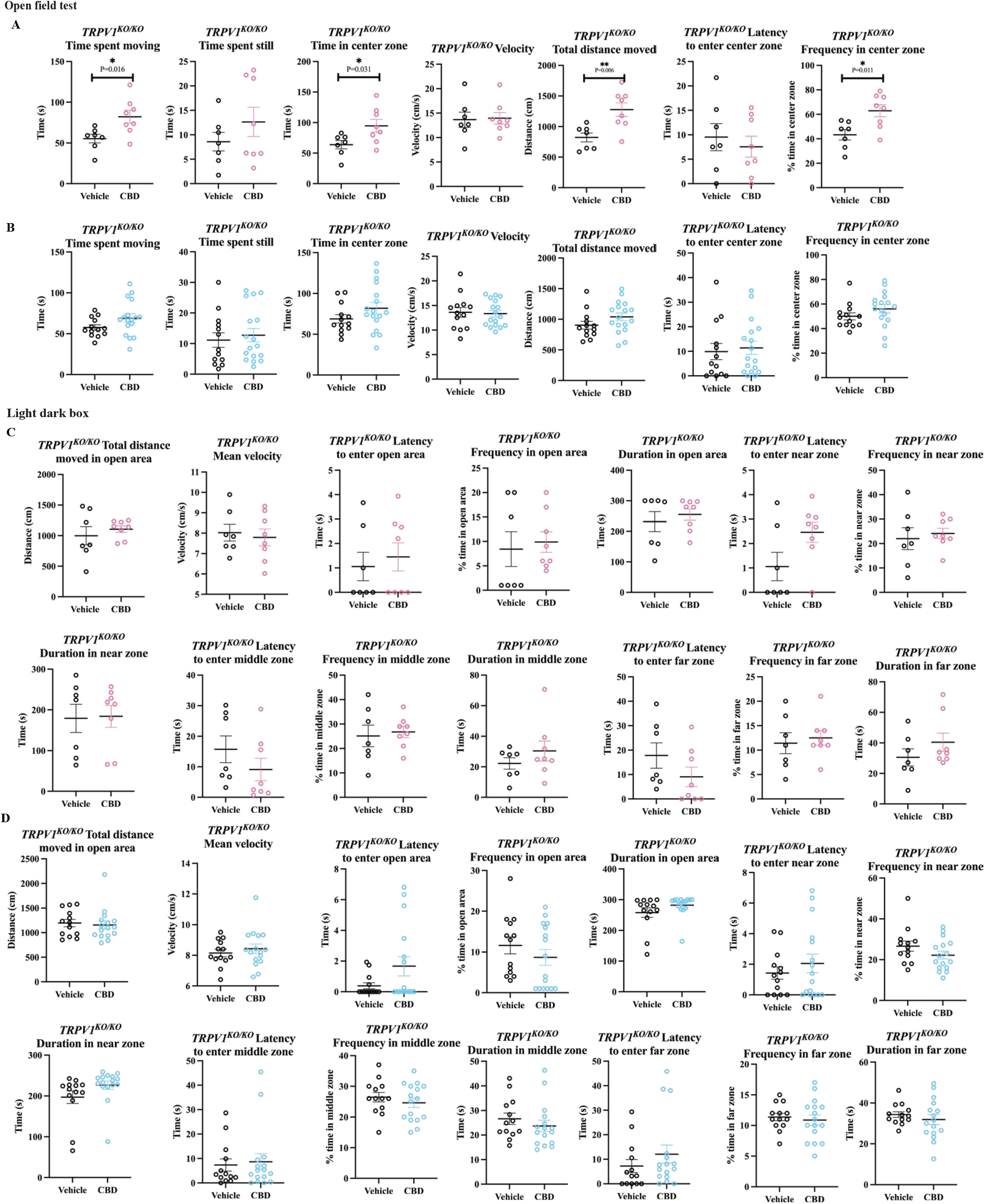
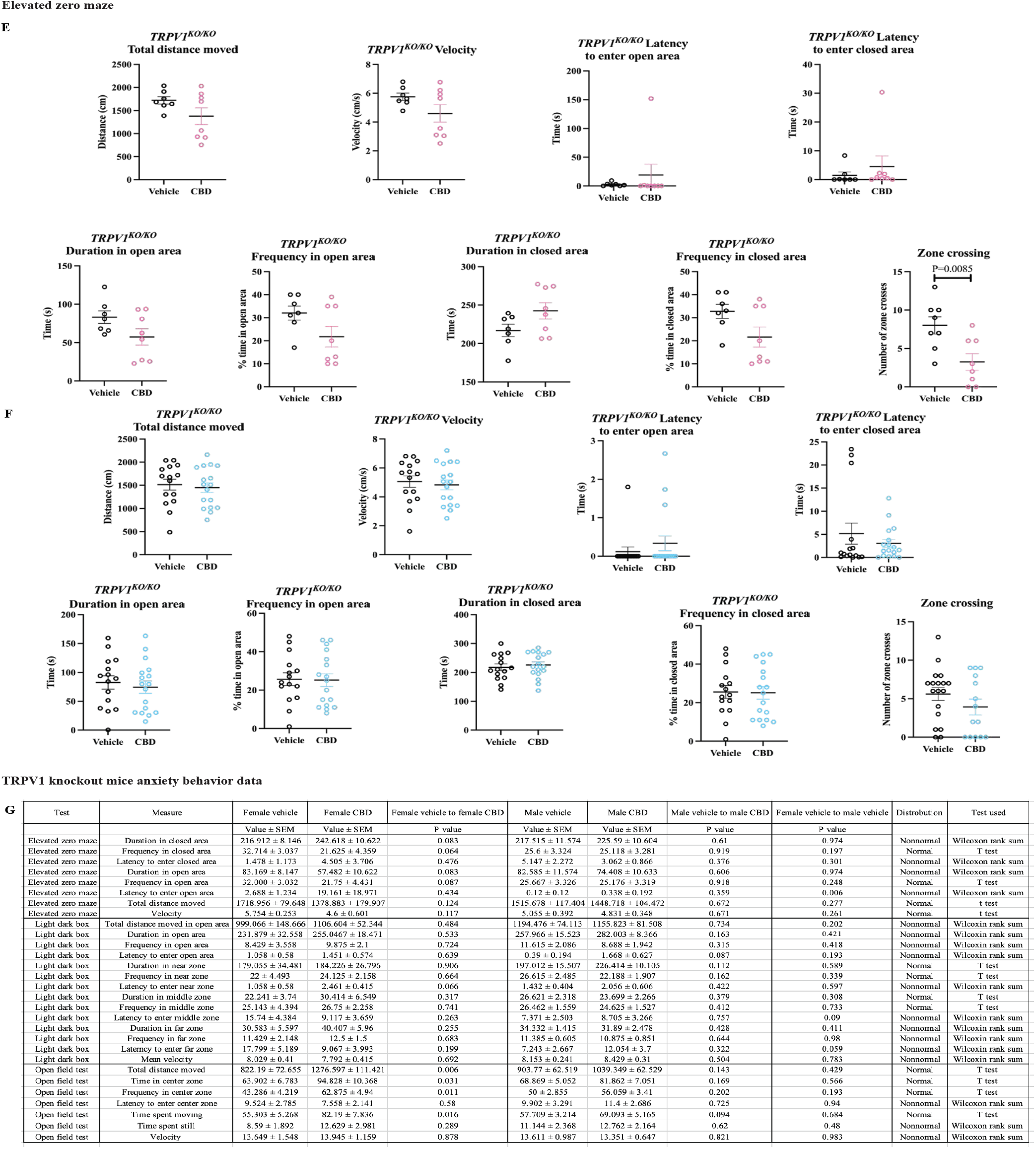
TRPV1 anxiety measures. **Fetal CBD exposure, overall, does not affect *TRPV1*^*KO/KO*^ anxiety.** Graphs show that fetal CBD exposure for *TRPV1*^*KO/KO*^ female offspring increases time spent moving (55.503 ± 5.268 seconds vehicle female, 82.19 ± 7.836 seconds, P=0.016, t-test), does not affect time spent still, increases time in center zone 63.902 ± 6.783 vehicle, 94.828 ± 10.368, P=0.031, t-test), does not affect velocity, increases total distance moved 822.19 ± 72.66 cm vehicle, 1276.60 ± 111.421 cm CBD, P=0.006, t-test), does not affect latency to enter center zone, and increases frequency in center zone via the open field test (43.286 ± 4.219 vehicle, 62.875 ± 4.94, P=0.011, t-test) (A). Graphs show that fetal CBD exposure for *TRPV1*^*KO/KO*^ male offspring does not alter time spent moving, time spent still, time in center zone, velocity, total distance moved, latency to enter center zone, and frequency in center zone via the open field test (B). Fetal CBD exposure does not affect female (C) or male (D) *TRPV1*^*KO/KO*^ offspring total distance moved in open area, mean velocity, latency to enter open area, frequency in open area, duration in open area, latency to enter near zone, frequency in near zone, latency to enter middle zone, frequency in middle zone, duration in middle zone, latency to enter far zone, frequency in far zone, or duration in far zone in the light dark box. Fetal CBD exposure does not affect female (E), or male (F) *TRPV1*^*KO/KO*^ offspring total distance moved, velocity, latency to enter open area, latency to enter closed area, duration in open area, frequency in open area, duration in closed area, or frequency in closed area in the elevated zero maze test. Fetal CBD exposure decreased the number of zone crosses of female *TRPV1*^*KO/KO*^ offspring (8.000 ± 3.162 vehicle, 3.25 ± 3.059 CBD, P=0.009, t-test). Table shows statistical analyses of all TRPV1KO vehicle-exposed and CBD-exposed measures of anxiety (G).

## Acknowledgements

We thank all members of the Bates lab for comments on the manuscript. We thank Dr. Nicolas Busquet from the Animal Behavior Core for essential advice on behavior experimental planning and for training us on the behavior apparatuses. We thank Dr. Cristina Sempio and Dr. Jost Klawitter for their advisement on CBD pharmacokinetics and metabolite analyses. We thank Dr. Erica Wymore and Dr. David Kroll for their guidance and expertise on the clinical and pharmacologic aspects of CBD consumption during pregnancy. This work was supported by the Institute of Cannabis Research, the Colorado Clinical and Translational Sciences Institute (CTSA Grant TL1 TR002533), and the University of Colorado Anschutz Medical Campus Diabetes Research Center.

## Author contributions

Conceptualization, Methodology, Formal Analysis, Investigation, Data Curation, Resources; K.S.S. and E.A.B. Writing original draft, review and editing, visualization: K.S.S. and E.A.B. Supervision: E.A.B. Project administration: E.A.B. Funding Acquisition K.S.S. and E.A.B. Pyramidal neuron analyses, writing original draft, review, editing, and visualization: L.E.G.W., V.H., W.C.O. Hargreaves tests; L.F. and E.A.B. Marble burying data analysis, K.K.

## Declaration of interests

The authors declare no competing interests.

## Notes

The authors declare no conflicts of interest.

### Competing Interest Statement

The authors have declared no competing interest.

