## Supplemental Table 1 and Supplemental Figures for "Fetal cannabidiol (CBD) exposure alters thermal pain sensitivity, cognition, and prefrontal cortex excitability"

**Supplemental table 1. Summary of anxiety measure statistics.** Table shows all measures for the open field test, light dark box, and elevated zero maze split by sex and statistical analysis.

**Supplemental table 1. Data for anxiety tests**

| Test | Measure | Female Vehicle | Female CBD | Female vehicle to female CBD | Male Vehicle | Male CBD | Male vehicle to Male CBD | Distrobution | Test used |
|---|---|---|---|---|---|---|---|---|---|
| | | Value ± SEM | Value ± SEM | P value | Value ± SEM | Value ± SEM | P value | | |
| Open field test | Freuqnecy in center zone | 49.93 ± 3.83 | 50.12 ± 2.29 | 0.965 | 49.26 ± 4.78 | 48.87 ± 2.88 | 0.942 | Normal | T-Test |
| Open field test | Velocity | 11.34 ± 0.92 | 10.88 ± 0.51 | 0.525 | 10.69 ± 0.63 | 10.71 ± 0.54 | 0.978 | Nonnormal | Wilcoxin rank sum test |
| Open field test | Total distance moved | 932.47 ± 81.25 | 925.19 ± 50.00 | 0.936 | 913.97 ± 90.52 | 930.76 ± 53.62 | 0.869 | Normal | T-Test |
| Open field test | Time in center zone | 90.20 ± 11.05 | 89.19 ± 6.36 | 0.933 | 89.40 ± 10.05 | 90.76 ± 6.00 | 0.904 | Normal | T-Test |
| Open field test | Time moving | 70.46 ± 7.65 | 71.90 ± 4.62 | 0.865 | 70.76 ± 6.95 | 72.53 ± 4.07 | 0.821 | Normal | T-Test |
| Open field test | Time still | 19.72 ± 3.82 | 17.28 ± 2.10 | 0.225 | 18.63 ± 3.72 | 18.23 ± 2.35 | 0.551 | Nonnormal | Wilcoxin rank sum test |
| Light dark box | Total distance moved in open area | 740.45 ± 106.50 | 1033.55 ± 92.01 | 0.071 | 854.22 ± 89.37 | 1191.09 ± 165.01 | 0.105 | Nonnormal | Wilcoxin rank sum test |
| Light dark box | Mean velocity | 6.79 ± 0.30 | 6.83 ± 0.29 | 0.572 | 7.12 ± 0.23 | 6.99 ± 0.50 | 0.484 | Nonnormal | Wilcoxin rank sum test |
| Light dark box | Duration in open area | 263.09 ± 17.52 | 234.12 ± 11.89 | 0.225 | 233.82 ± 18.18 | 246.27 ± 14.30 | 0.239 | Nonnormal | Wilcoxin rank sum test |
| Light dark box | Duration in near zone | 219.84 ± 16.48 | 159.26 ± 15.01 | 0.052 | 180.42 ± 19.22 | 179.64 ± 13.14 | 0.973 | Normal | T-Test |
| Light dark box | Duration in middle zone | 17.34 ± 3.37 | 27.23 ± 2.54 | 0.082 | 21.65 ± 2.99 | 31.87 ± 2.38 | 0.014 | Normal | T-Test |
| Light dark box | Duration in far zone | 25.28 ± 5.92 | 47.62 ± 11.22 | 0.306 | 31.75 ± 3.49 | 37.65 ± 2.78 | 0.156 | Nonnormal | Wilcoxin rank sum test |
| Light dark box | Latency to enter near zone | 1.49 ± 0.82 | 6.90 ± 4.79 | 0.978 | 3.57 ± 2.82 | 3.61 ± 1.68 | 0.766 | Nonnormal | Wilcoxin rank sum test |
| Light dark box | Latency to enter middle zone | 2.15 ± 1.17 | 18.85 ± 6.12 | 0.112 | 12.98 ± 4.98 | 13.35 ± 4.37 | 0.915 | Nonnormal | Wilcoxin rank sum test |
| Light dark box | Latency to enter far zone | 10.488 ± 5.497 | 20.30 ± 6.25 | 0.428 | 16.19 ± 5.65 | 15.85 ± 4.60 | 0.965 | Nonnormal | Wilcoxin rank sum test |
| Light dark box | Frequency in near zone | 19.83 ± 3.84 | 27.33 ± 2.87 | 0.207 | 21.23 ± 2.52 | 24.79 ± 2.67 | 0.391 | Normal | T-Test |
| Light dark box | Frequency in middle zone | 5.59 ± 2.28 | 20.33 ± 1.97 | 0.184 | 18.62 ± 1.95 | 22.25 ± 1.72 | 0.196 | Normal | T-Test |
| Light dark box | Frequency in far zone | 7.00 ± 1.39 | 10.95 ± 1.55 | 0.124 | 8.85 ± 0.89 | 10.54 ± 0.87 | 0.193 | Nonnormal | Wilcoxin rank sum test |
| Elevated zero maze | Velocity | 5.27 ± 0.36 | 5.25 ± 0.26 | 0.969 | 4.79 ± 0.26 | 4.92 ± 0.26 | 0.733 | Normal | T-Test |
| Elevated zero maze | Frequency in light area | 33.85 ± 3.35 | 31.68 ± 2.17 | 0.577 | 27.43 ± 2.15 | 27.04 ± 2.29 | 0.901 | Normal | T-Test |
| Elevated zero maze | Total distance moved | 1573.94 ± 108.78 | 1570.05 ± 77.47 | 0.976 | 1437.06 ± 78.82 | 1474.54 ± 79.04 | 0.739 | Normal | T-Test |
| Elevated zero maze | Time in closed area | 212.67 ± 9.16 | 206.69 ± 6.93 | 0.977 | 222.98 ± 6.21 | 218.35 ± 5.76 | 0.395 | Nonnormal. | Wilcoxin rank sum test |
| Elevated zero maze | Time in open area | 87.43 ± 9.16 | 93.41 ± 6.93 | 0.977 | 77.12 ± 6.22 | 81.75 ± 5.76 | 0.395 | Nonnormal | Wilcoxin rank sum test |
| Elevated zero maze | Frequncy in closed area | 34 ± 3.34 | 31.6 ± 2.14 | 0.534 | 27.30 ± 2.18 | 26.92 ± 2.32 | 0.904 | Normal | T-Test |

**Supplemental Figure 1. Fetal CBD exposure, overall, does not affect *TRPV1$^{KO/KO}$* anxiety.** Graphs show that fetal CBD exposure for *TRPV1$^{KO/KO}$* female offspring increases time spent moving (55.503 ± 5.268 seconds vehicle female, 82.19 ± 7.836 seconds, P=0.016, t-test), does not affect time spent still, increases time in center zone 63.902 ± 6.783 vehicle, 94.828 ± 10.368, P=0.031, t-test), does not affect velocity, increases total distance moved 822.19 ± 72.66 cm vehicle, 1276.60 ± 111.421 cm CBD, P=0.006, t-test), does not affect latency to enter center zone, and increases frequency in center zone via the open field test (43.286 ± 4.219 vehicle, 62.875 ± 4.94, P=0.011, t-test) (A). Graphs show that fetal CBD exposure for *TRPV1$^{KO/KO}$* male offspring does not alter time spent moving, time spent still, time in center zone, velocity, total distance moved, latency to enter center zone, and frequency in center zone via the open field test (B). Fetal CBD exposure does not affect female (C) or male (D) *TRPV1$^{KO/KO}$* offspring total distance moved in open area, mean velocity, latency to enter open area, frequency in open area, duration in open area, latency to enter near zone, frequency in near zone, latency to enter middle zone, frequency in middle zone, duration in middle zone, latency to enter far zone, frequency in far zone, or duration in far zone in the light dark box. Fetal CBD exposure does not affect female (E), or male (F) *TRPV1*$^{KO/KO}$ offspring total distance moved, velocity, latency to enter open area, latency to enter closed area, duration in open area, frequency in open area, duration in closed area, or frequency in closed area in the elevated zero maze test. Fetal CBD exposure decreased the number of zone crosses of female *TRPV1*$^{KO/KO}$ offspring (8.000 ± 3.162 vehicle, 3.25 ± 3.059 CBD, P=0.009, t-test). Table shows statistical analyses of all TRPV1KO vehicle-exposed and CBD-exposed measures of anxiety (G).

**Supplemental figure 1. TRPV1 anxiety measures**

Open field test

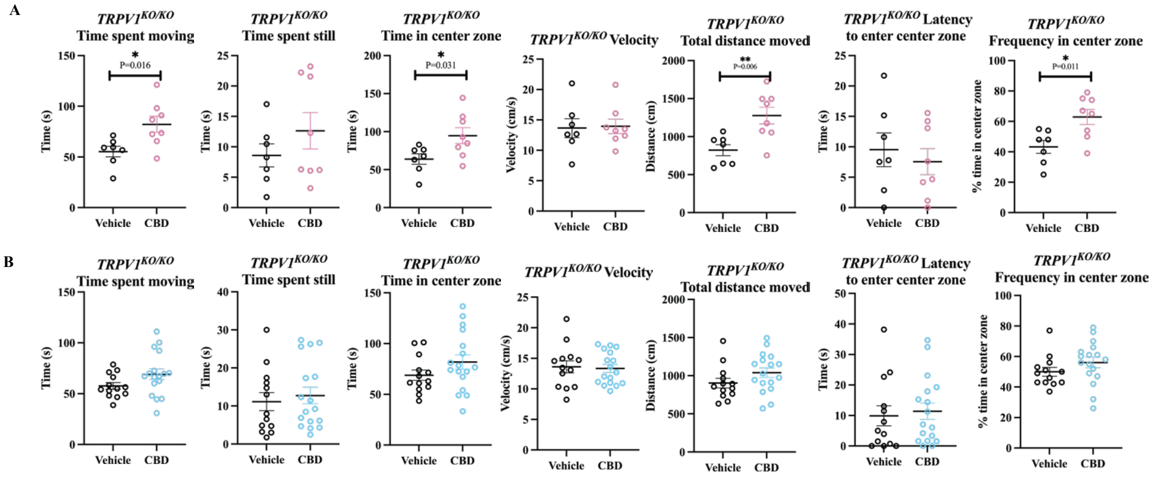


Light dark box

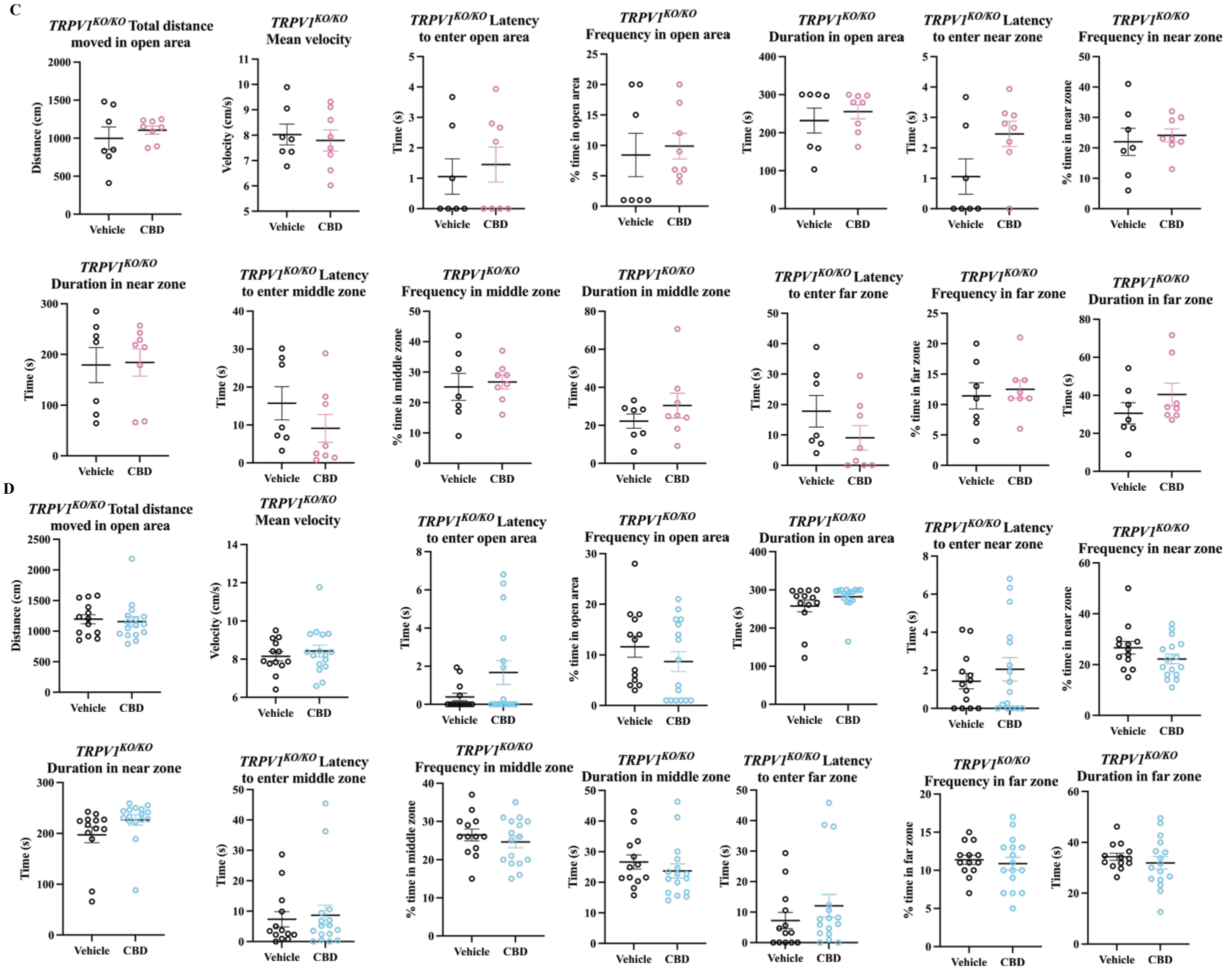

Elevated zero maze

E

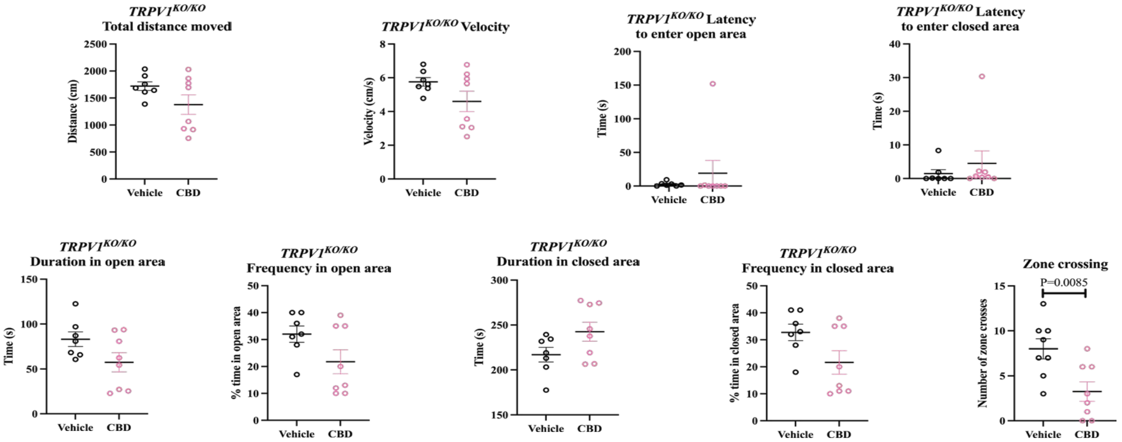


F

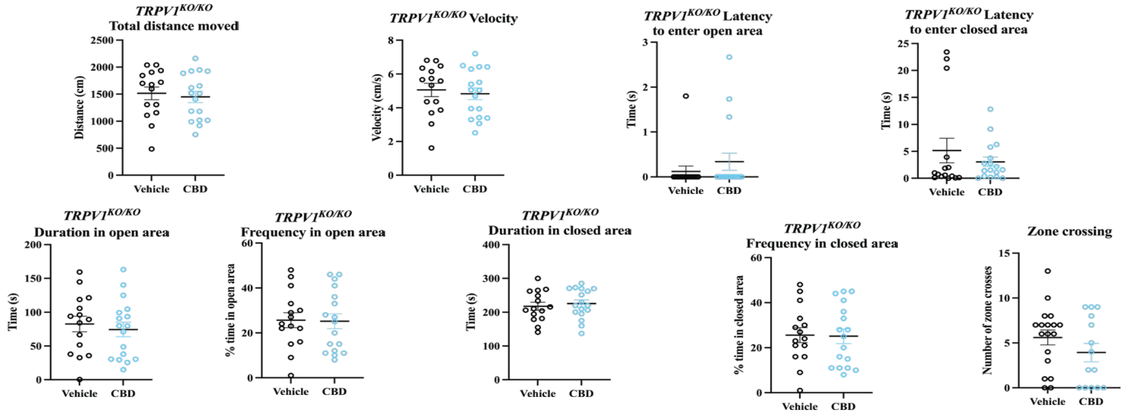


TRPV1 knockout mice anxiety behavior data

G

| Test | Measure | Female vehicle | Female CBD | Female vehicle to female CBD | Male vehicle | Male CBD | Male vehicle to male CBD | Female vehicle to male vehicle | Distrobution | Test used |
|---|---|---|---|---|---|---|---|---|---|---|
| | | Value ± SEM | Value ± SEM | P value | Value ± SEM | Value ± SEM | P value | P value | | |
| Elevated zero maze | Duration in closed area | 216.912 ± 8.146 | 242.618 ± 10.622 | 0.083 | 217.515 ± 11.574 | 225.59 ± 10.604 | 0.61 | 0.974 | Nonnormal | Wilcoxon rank sum |
| Elevated zero maze | Frequency in closed area | 32.714 ± 3.037 | 21.625 ± 4.359 | 0.064 | 25.6 ± 3.324 | 25.118 ± 3.281 | 0.919 | 0.197 | Normal | T test |
| Elevated zero maze | Latency to enter closed area | 1.478 ± 1.173 | 4.505 ± 3.706 | 0.476 | 5.147 ± 2.272 | 3.062 ± 0.866 | 0.376 | 0.301 | Nonnormal | Wilcoxon rank sum |
| Elevated zero maze | Duration in open area | 83.169 ± 8.147 | 57.482 ± 10.622 | 0.083 | 82.585 ± 11.574 | 74.408 ± 10.633 | 0.606 | 0.974 | Nonnormal | Wilcoxon rank sum |
| Elevated zero maze | Frequency in open area | 32.000 ± 3.032 | 21.75 ± 4.431 | 0.087 | 25.667 ± 3.326 | 25.176 ± 3.319 | 0.918 | 0.248 | Normal | T test |
| Elevated zero maze | Latency to enter open area | 2.688 ± 1.234 | 19.161 ± 18.971 | 0.434 | 0.12 ± 0.12 | 0.338 ± 0.192 | 0.359 | 0.006 | Nonnormal | Wilcoxon rank sum |
| Elevated zero maze | Total distance moved | 1718.956 ± 79.648 | 1378.883 ± 179.907 | 0.124 | 1515.678 ± 117.404 | 1448.718 ± 104.472 | 0.672 | 0.277 | Normal | t test |
| Elevated zero maze | Velocity | 5.754 ± 0.253 | 4.6 ± 0.601 | 0.117 | 5.055 ± 0.392 | 4.831 ± 0.348 | 0.671 | 0.261 | Normal | t test |
| Light dark box | Total distance moved in open area | 999.066 ± 148.666 | 1106.604 ± 52.344 | 0.484 | 1194.476 ± 74.113 | 1155.823 ± 81.508 | 0.734 | 0.202 | Nonnormal | Wilcoxin rank sum |
| Light dark box | Duration in open area | 231.879 ± 32.558 | 255.0467 ± 18.471 | 0.533 | 257.966 ± 15.523 | 282.003 ± 8.366 | 0.163 | 0.421 | Nonnormal | Wilcoxin rank sum |
| Light dark box | Frequency in open area | 8.429 ± 3.558 | 9.875 ± 2.1 | 0.724 | 11.615 ± 2.086 | 8.688 ± 1.942 | 0.315 | 0.418 | Nonnormal | Wilcoxin rank sum |
| Light dark box | Latency to enter open area | 1.058 ± 0.58 | 1.451 ± 0.574 | 0.639 | 0.39 ± 0.194 | 1.668 ± 0.627 | 0.087 | 0.193 | Nonnormal | Wilcoxin rank sum |
| Light dark box | Duration in near zone | 179.055 ± 34.481 | 184.226 ± 26.796 | 0.906 | 197.012 ± 15.507 | 226.414 ± 10.105 | 0.112 | 0.589 | Normal | T test |
| Light dark box | Frequency in near zone | 22 ± 4.493 | 24.125 ± 2.158 | 0.664 | 26.615 ± 2.485 | 22.188 ± 1.907 | 0.162 | 0.339 | Normal | T test |
| Light dark box | Latency to enter near zone | 1.058 ± 0.58 | 2.461 ± 0.415 | 0.066 | 1.432 ± 0.404 | 2.056 ± 0.606 | 0.422 | 0.597 | Nonnormal | Wilcoxin rank sum |
| Light dark box | Duration in middle zone | 22.241 ± 3.74 | 30.414 ± 6.549 | 0.317 | 26.621 ± 2.318 | 23.699 ± 2.266 | 0.379 | 0.308 | Normal | T test |
| Light dark box | Frequency in middle zone | 25.143 ± 4.394 | 26.75 ± 2.258 | 0.741 | 26.462 ± 1.559 | 24.625 ± 1.527 | 0.412 | 0.733 | Normal | T test |
| Light dark box | Latency to enter middle zone | 15.74 ± 4.384 | 9.117 ± 3.659 | 0.263 | 7.371 ± 2.503 | 8.705 ± 3.266 | 0.757 | 0.09 | Nonnormal | Wilcoxin rank sum |
| Light dark box | Duration in far zone | 30.583 ± 5.597 | 40.407 ± 5.96 | 0.255 | 34.332 ± 1.415 | 31.89 ± 2.478 | 0.428 | 0.411 | Nonnormal | Wilcoxin rank sum |
| Light dark box | Frequency in far zone | 11.429 ± 2.148 | 12.5 ± 1.5 | 0.683 | 11.385 ± 0.605 | 10.875 ± 0.851 | 0.644 | 0.98 | Nonnormal | Wilcoxin rank sum |
| Light dark box | Laterncy to enter far zone | 17.799 ± 5.189 | 9.067 ± 3.993 | 0.199 | 7.243 ± 2.667 | 12.054 ± 3.7 | 0.322 | 0.059 | Nonnormal | Wilcoxin rank sum |
| Light dark box | Mean velocity | 8.029 ± 0.41 | 7.792 ± 0.415 | 0.692 | 8.153 ± 0.241 | 8.429 ± 0.31 | 0.504 | 0.783 | Nonnormal | Wilcoxin rank sum |
| Open field test | Total distance moved | 822.19 ± 72.655 | 1276.597 ± 111.421 | 0.006 | 903.77 ± 62.519 | 1039.349 ± 62.529 | 0.143 | 0.429 | Normal | T test |
| Open field test | Time in center zone | 63.902 ± 6.783 | 94.828 ± 10.368 | 0.031 | 68.869 ± 5.052 | 81.862 ± 7.051 | 0.169 | 0.566 | Normal | T test |
| Open field test | Frequency in center zone | 43.286 ± 4.219 | 62.875 ± 4.94 | 0.011 | 50 ± 2.855 | 56.059 ± 3.41 | 0.202 | 0.193 | Normal | T test |
| Open field test | Latency to enter center zone | 9.524 ± 2.785 | 7.558 ± 2.141 | 0.58 | 9.902 ± 3.291 | 11.4 ± 2.686 | 0.725 | 0.94 | Nonnormal | Wilcoxon rank sum |
| Open field test | Time spent moving | 55.303 ± 5.268 | 82.19 ± 7.836 | 0.016 | 57.709 ± 3.214 | 69.093 ± 5.165 | 0.094 | 0.684 | Normal | T test |
| Open field test | Time spent still | 8.59 ± 1.892 | 12.629 ± 2.981 | 0.289 | 11.144 ± 2.368 | 12.762 ± 2.164 | 0.62 | 0.48 | Nonnormal | Wilcoxon rank sum |
| Open field test | Velocity | 13.649 ± 1.548 | 13.945 ± 1.159 | 0.878 | 13.611 ± 0.987 | 13.351 ± 0.647 | 0.821 | 0.983 | Nonnormal | Wilcoxon rank sum |
